# Dynamic metabolic reprogramming in dendritic cells: an early response to influenza infection that is essential for effector function

**DOI:** 10.1101/2020.01.14.906826

**Authors:** Svetlana Rezinciuc, Lavanya Bezavada, Azadeh Bahadoran, Susu Duan, Ruoning Wang, Daniel Lopez-Ferrer, Erika E. Zink, David Finklestein, Douglas R. Green, Ljiljana Pasa-Tolic, Paul G. Thomas, Heather S. Smallwood

**Author notes:** Corresponding author (HS).

## Abstract

Infection with the influenza virus triggers an innate immune response aimed at initiating the adaptive response to halt viral replication and spread. However, the metabolic response fueling the molecular mechanisms underlying changes in innate immune cell homeostasis remain undefined. Thus, we compared the metabolic response of dendritic cells to that of those infected with active and inactive influenza A virus or treated with toll like receptor agonists. While influenza infects dendritic cells, it does not productively replicate in these cells, and therefore metabolic changes upon infection may represent an adaptive response on the part of the host cells. Using quantitative mass spectrometry along with pulse chase substrate utilization assays and metabolic flux measurements, we found global metabolic changes 17 hours post infection, including significant changes in carbon commitment via glycolysis and glutaminolysis, as well as ATP production via TCA cycle and oxidative phosphorylation. Influenza infection of dendritic cells led to a metabolic phenotype, distinct from that induced by TLR agonists, with significant resilience in terms of metabolic plasticity. We identified Myc as one transcription factor modulating this response. Restriction of either Myc activity or mitochondrial substrates resulted in significant changes in the innate immune functions of dendritic cells, including reduced motility and T cell activation. Transcriptome analysis of inflammatory dendritic cells isolated following influenza infection showed similar metabolic reprogramming occurs in vivo. Thus, early in the infection process dendritic cells respond with global metabolic restructuring that is present in lung DC 9 days following infection and impacts their effector function, suggesting that metabolic switching in dendritic cells plays a vital role in initiating the immune response to influenza infection.

**Author Summary:** In response to influenza infection we found that dendritic cells, cells that are critical in mounting an effective immune response, undergo a profound metabolic shift. They alter the concentration and location of hundreds of proteins, including c-MYC, mediating a shift to a highly glycolytic phenotype that is also flexible in terms of fueling respiration. Dendritic cells initiate the immune response to influenza and activate the adaptive response allowing viral clearance and manifesting immune memory for protection against subsequent infections. We found that limiting access to specific metabolic pathways or substrates diminished key immune functions. Previously we described an immediate, fixed, hypermetabolic state in infected respiratory epithelial cells. We now show the metabolic responses of epithelial and dendritic cells are distinct. Here, we also demonstrate that dendritic cells tailor their metabolic response to the pathogen or TLR stimulus. This metabolic reprogramming occurs rapidly in vitro and it is sustained in inflammatory dendritic cells in vivo for at least 9 days following influenza infection. Thus, drugs targeting metabolism are likely to have cell- and pathogen-specific activities in the context of infection. These studies open the possibility of modulating the immune response to viral infection via customizing metabolic therapy to enhance or diminish the function of specific cells.

## Introduction

The influenza virus is associated with significant disease burden in the human population and is of particularly high risk to children, the elderly, and those with certain medical conditions, such as pregnancy, obesity, or metabolic disease. In a productive infection, the influenza virus co-opts the host cell’s molecular machinery to induce a virus centric shift in macromolecular production for budding virions. In connection to this, we have previously shown influenza A virus infection of human respiratory epithelial cells leads to a fixed hypermetabolic state[1]. Dendritic cells are dispersed throughout the respiratory tract (including the lung parenchyma, alveolar space, and the airway epithelium), are a heterogeneous population with respect to surface phenotype and function, and are constantly surveying and sampling for pathogens or foreign material at this delicate barrier between the environment and blood stream [2–5]. Following IAV infection, dendritic cells have a critical role linking the innate and adaptive immune responses. These antigen presenting cells are dependent on the binding of different pattern-recognition receptors binding to pathogen-associated molecules such as toll-like receptors (TLRs), typically engaged on the cell surface or in endosomes. Influenza is sensed intracellularly by TLR3, TLR7, TLR8, and TLR9, sensing either viral envelope proteins or nucleic acids, respectively [6–12]. These TLRs are expressed at variable levels in human and murine cells of the upper respiratory tract, such as epithelial cells, macrophages, and dendritic cells [13]. Combined stimulation of these intracellular receptors tailor the antiviral response via retinoic acid-inducible gene-I (RIG-I) and NOD-like receptors, or lack thereof, creating a cell-specific response to influenza infection [13]. One molecular driver underpinning immune activation is an increase in metabolism to fuel effector functions requisite for pathogen elimination [14–17]. Recently, seminal papers in the field have demonstrated that activation of dendritic cells with the TLR4 agonist lipopolysaccharide (LPS) has profound metabolic consequences that may correlate to altered immune function [17–19]. However, the metabolic response of dendritic cells to influenza infection remains undefined, as does its relationship to TLR stimulation and inactive virus.

To gain a better understanding of the molecular events that initiate the systemic immune response to influenza infection, we performed quantitative proteomics on influenza A virus strain A/Puerto Rico/8/1934 H1N1 (IAV) infected and uninfected bone marrow derived dendritic cells (DC) that were differentiated in vitro. Given that cellular location is pivotal to metabolic enzyme activity, we divided the DC proteins into soluble and insoluble fractions to increase proteome coverage while defining protein redistribution in response to infection. We found that metabolic pathways dominated the enriched protein networks, accompanied by localization changes in metabolic enzymes in response to IAV infection. The functional impact of these abundance and localization changes were determined with pulse chase experiments using labeled metabolic substrates, bioenergetics, and metabolite quantification. Similar to our previous findings in epithelial cells[1], we identified upregulation of the transcription factor c-MYC and found that c-MYC inhibition restored infected DC bioenergetics to quiescent levels. However, unlike IAV-infected epithelial cells, infected DC increased their metabolic plasticity by reprogramming glycolysis, glutaminolysis, and respiration better adapting to use diverse fuels to meet the challenges of high energy demand and anabolism needed for activation. To dissect the DC response to specific activation pathways, we quantified metabolic flux rates in response to TLR3, 4, or 7/8 agonists, genome replication-incompetent virus, and wild type virus. Intriguingly, DC displayed treatment specific differences in glycolysis and mitochondrial respiration concomitant to differences in coupling efficiency and ATP production. The response of DC to viral infection induced a consistent and robust increase in glycolysis that was decoupled from the canonical TCA cycle. Our results revealed that IAV-infected DC rely on glycolysis and glutamine over pyruvate utilization. Further, we analyzed changes in infected DC motility, phagocytosis, and T cell activation and found that inhibiting c-MYC activity or import of specific mitochondrial substrates significantly reduced key immune functions. We then determined that this metabolic reprogramming occurs in inflammatory lung dendritic cells 9 days following infection and results in a very significant increase in glycolytic enzymes over those in the TCA cycle. Thus, in response to IAV infection, global protein expression and localization changes occur in DC and are associated with controlled metabolic restructuring that is necessary for DC function. Consequently, loss of metabolic homeostasis in dendritic cells or their lung microenvironment may have profound functional consequence for the overall immune response to influenza infection.

## Results

### Influenza induces significant changes in DC metabolic proteins and pathways

Two quantitative labeling strategies were used to determine proteomic changes after influenza virus infection. We infected in vitro differentiated DC with the PR8 mouse-adapted H1N1 influenza strain A/Puerto Rico/68/34 (IAV) at a multiplicity-of-infection (MOI) of 5 for 17 hours. In mouse DC, IAV replication is abortive (i.e. IAV replicates and virions build up intracellularly instead of being released) and has a much lower infectivity rate than in human DC [20–22]. Thus, we used a higher MOI than in our previous studies and validated effective infection of DC at this time point, as determined by viral protein production (S1A Fig). After cells were harvested, soluble and insoluble fractions were subjected to specialized trypsinization strategies to enhance coverage of proteins with complex tertiary structures (i.e., either filter-aided sample preparation (FASP) digestion, isobaric tags for relative and absolute quantitative (iTRAQ) labeling, or several rounds of high intensity focused ultrasound (HIFU) and pressurization followed by stable isotope labeling (SIL) (S1B Fig). The entire DC proteome (soluble and insoluble fractions ± infection) are publicly available at the Proteomics Identifications (PRIDE) repository (PRIDE access number made available on acceptance for publication). Approximately 25,000 peptides were detected with corresponding reporter ions; of these, 7520 mapped to National Center for Biotechnology Information (NCBI) reference sequence accession numbers (Table A S1 File). Of the 7,520 peptides, 70% had 2 or more peptide hits and were included in the quantitative analysis: 72% of -fold change in peptide abundance (Table 1). Most dynamic changes occurred in the insoluble fraction, with 855 proteins increased and 649 decreased after IAV infection (Table 1 and S1C Fig).

**Table 1.** Net changes in DC proteins in response to IAV infection after 17 hours. iTRAQ and SIL analysis of influenza virus infected BMDC. The intensities of the isobaric tag reporter ions were quantified by using the MASIC tool with the exclusion of missing reporter-ion channels or by calculating the SIL ratio for each peptide pair after accounting for singly or doubly labeled species in the ^16^O/^18^O ratio and correcting for labeling efficiency. Then, the MS/MS data were searched and filtered by using 0.5% FDR; peptides passing the filter were quantified then peptides-to-protein rollup was performed.

| Fraction | Up | No Change | Down | Total |
| --- | --- | --- | --- | --- |
| **Insoluble** | **855** | **2547** | **649** | **4051** |
| **Soluble** | **239** | **2003** | **427** | **2669** |
| **Both** | **74** | **1103** | **64** | **1241** |

To validate the iTRAQ results, we used the well-established SIL method (^16^O/^18^O labeling) and HIFU with the same infection protocol [23]. This technique identified more than 20,000 peptides, resulting in about 10,000 unique International Protein Index accession numbers, and 2000 protein identifications. As in the iTRAQ DC proteomes, the majority (73%) of DC peptides remained unaltered by IAV infection, which in -fold change in the abundance of 27% of DC proteins (Table A S1 File). Similar to the iTRAQ proteome, SIL samples had more changes in the insoluble fraction. Although iTRAQ provided expanded coverage of the DC proteome, more than half of all peptides identified by using SIL were also present in the iTRAQ data sets, with similar trends (S1C Fig). Next, we used bioinformatic tools to identify the major pathways altered by influenza. Proteomic data were segregated into increasing and decreasing sectors of 2-fold change. iTRAQ and SIL proteomes were combined, and the most significant peptides were selected. The lists of upregulated and downregulated proteins were submitted to the Database for Annotation, Visualization, and Integrated Discovery (DAVID) v6.8 and queried for enrichment groups [24]. 17 hours following IAV infection of DC, we found that metabolism accounts for the majority of KEGG Orthology (KO) Class processes and was significantly enriched in both the decreasing and increasing proteomes (Fig 1A & B respectively). Surprisingly, only a small fraction (8.6% up and 11% down) of the enriched pathways were in the immune system or infectious disease KEGG KO subclasses in the BRITE hierarchy (Table B S1 File).

**Figure 1.**
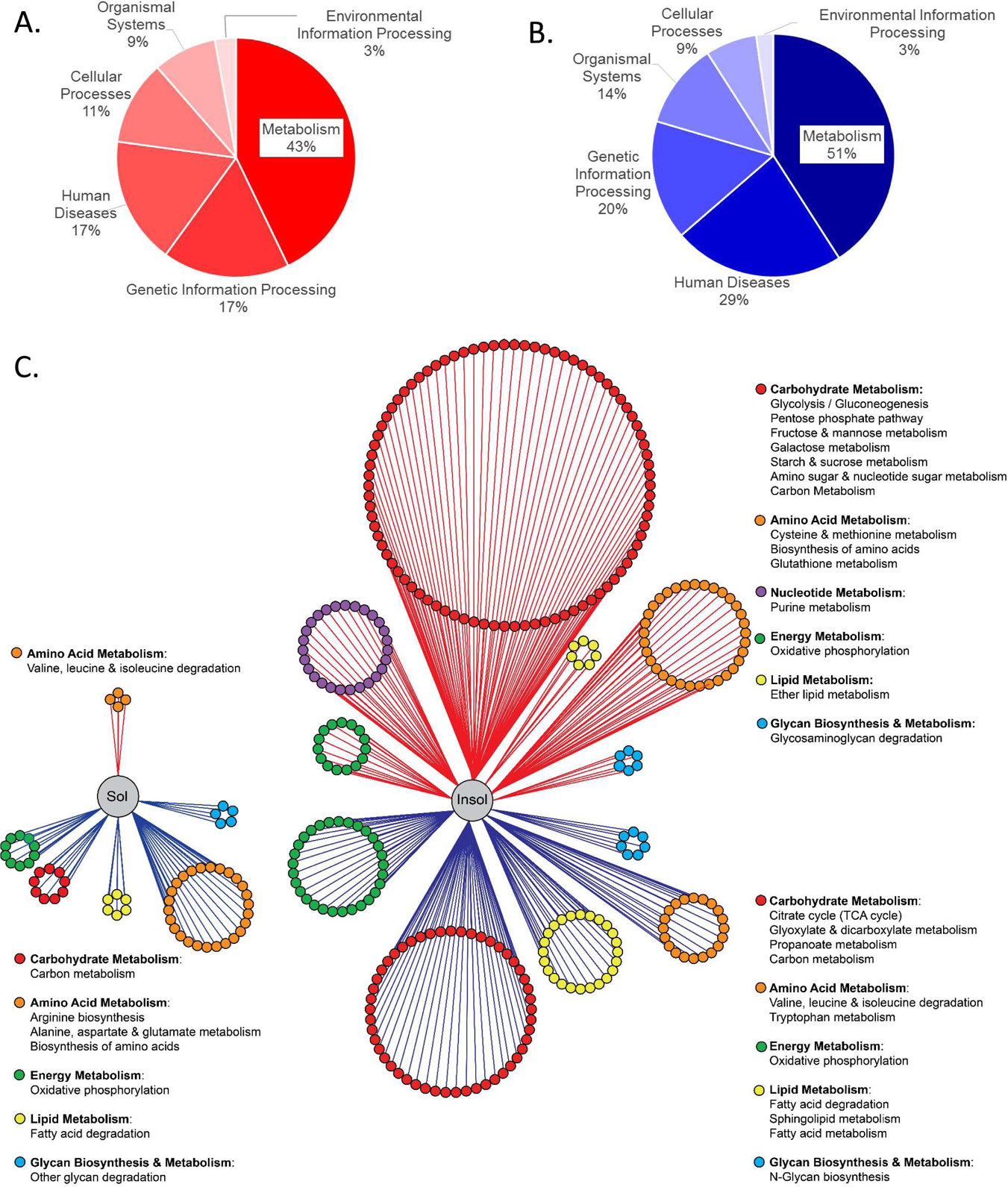
Infection induced changes in DC protein abundance and localization. Influenza virus strain A/PuertoRico/34 (IAV) was added to DC for 2 hours to allow entry, then medium replaced, and the infection proceeded for 17 hours followed by cell lysis and protein extraction. Proteins were labeled with either SIL or iTRAQ subjected to LC-MS/MS. Both proteomes were combined, redundancies removed and confidently identified peptides with abundance changes of 2-fold or greater linked to protein identifiers. The lists of upregulated and downregulated proteins were submitted to the Database for Annotation, Visualization, and Integrated Discovery (DAVID) v6.7 and mapped to KEGG pathways. (A-B) Significantly enriched KEGG pathways were grouped by KEGG Orthology (KO) Class for the increased (A) and decreased (B) proteomes. (C) In the network summary of metabolic protein dynamics, the soluble and insoluble proteomes were subdivided by abundance change. Proteins with relative expression changes of two-fold or greater are connected to their location in the soluble or insoluble fraction by red (top) or blue (bottom) edges indicating increased or decreased abundance change, respectively. Each protein node size was held static and color coded by KEGG subclass with the corresponding metabolic pathway the protein mapped to delineated in the legend. The nodes of each KEGG subclass were arranged into circles that are proportional to the number of proteins with abundance changes in each significantly enriched metabolic pathway.

A global view of influenza-induced changes in the abundance and localization of metabolic proteins is depicted in Figure 1C, represented as previously described [25]. In this network summary of metabolic protein dynamics, KO metabolic class proteins were organized based on location in the soluble or insoluble fraction (Fig 1C left and right, respectively). Metabolic proteins with relative expression changes of two-fold or greater were connected to their location in the soluble or insoluble fraction by red or blue edges (up or down, respectively). Each protein node is colored by KEGG subclass with significantly enriched KEGG metabolic pathways delineated in the legend. Protein coverage per pathway and statistical information for pathway enrichment is provided in Supplementary File 1 Tables B and C. Thus, the circle size of each KEGG metabolic class is proportional to the number of proteins with significant fold changes in it (Fig 1C). We found many of the proteins with significant abundance changes were involved in carbohydrate metabolism (Fig 1C red nodes and S2D Fig). In the insoluble increased proteome, we found carbohydrate metabolism had the largest number of proteins dispersed in several significantly enriched KEGG pathways such as glycolysis, pentose phosphate pathway, fructose/mannose, galactose, starch/sucrose, amino sugar/nucleotide metabolism, as well as carbon metabolism in general (Fig 1C red nodes with red edges). Indeed, in the increased proteome, carbohydrate metabolism accounted for 47% of the significantly enriched pathways, including glycolysis. On the other hand, in the insoluble decreased proteome, carbohydrate metabolism had fewer proteins, and these were associated with TCA cycle, glyoxylate/dicarboxylate, and propanoate metabolism (Fig 1C red nodes blue edges). This differential expression of proteins controlling carbon metabolism may indicate uncoupling of glycolysis from oxidative phosphorylation (OXPHOS).

We identified 432 differentially expressed metabolic proteins (i.e. <u>>2</u>-fold abundance changes) that mapped to well-known metabolic pathways (Table 2 and Table B S1 File). Significant enrichment of glycolysis in the KEGG analysis led us to more closely examine protein changes in this pathway. Coverage for glycolytic proteins was complete in both fractions. We found that most soluble glycolytic proteins in this network increased (S1C Fig green). The insoluble fraction had fewer proteins that increased two-fold or more, which is not surprising given that glycolysis occurs in the cytosol. Some proteins increased in both fractions, - -Enolase is a glycolytic protein that also rapidly increases and dimerizes at the surface of activated monocytes to form a plasminogen receptor critical for immune cell recruitment to the lung [26]. We validated-Enolase increase in both soluble and insoluble fractions with western blots (S1C Fig).

**Table 2.**
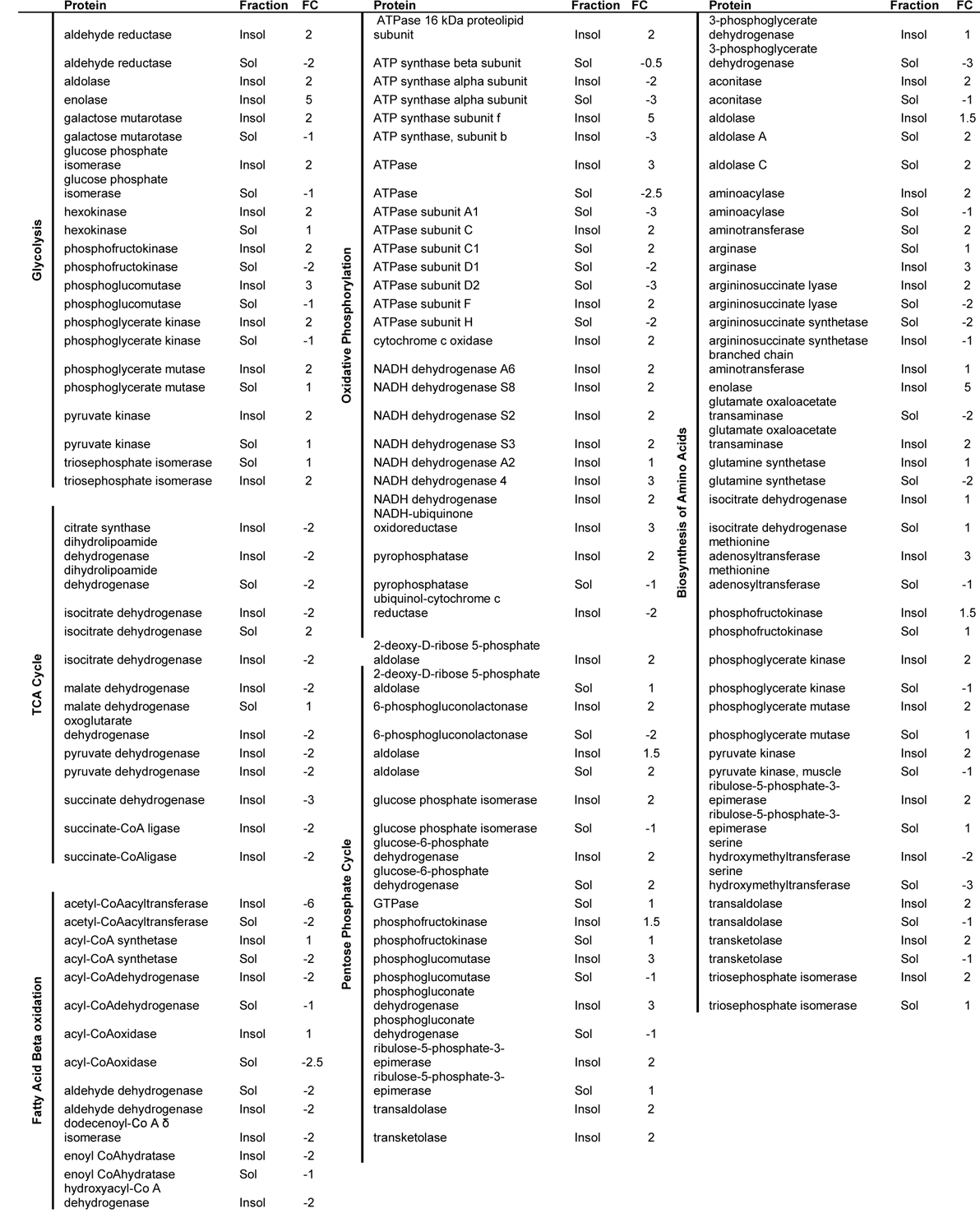
Infection induced changes in the relative abundance of DC metabolic proteins

Related to but not contingent upon expression levels, metabolic proteins are also regulated by changes in subcellular location [25, 27-30]. In response to infection, we detected 48 metabolic proteins that relocalized from the soluble to insoluble fraction or vice versa (Table 2). Some of the proteins undergoing localization changes were associated with metabolic pathways. For example, we found proteins involved in valine, leucine, and isoleucine degradation were decreased in the insoluble proteome while increased in the soluble (Fig 1C orange nodes and Table B & C S1 File). Conversely, we found key players in the biosynthesis of amino acids decreased in the soluble and increased in the insoluble fractions (Fig 1C orange nodes and Table B & C S1 File). We also found metabolic pathways that were uniformly altered irrespective of location such as the fatty acid degradation pathway (Fig 1C yellow nodes and Table C S1 File). Taken together, these results demonstrate that in response to IAV infection, DC coordinate changes in metabolic protein abundances and location.

Table 2. Infection induced changes in DC protein abundance and localization. Influenza virus strain A/PuertoRico/68/34 (IAV) was added to DC for 2 hours, infection medium replaced, and the infection proceeded for 17 hours followed by cell lysis and protein extraction. Proteins were labeled with either SIL or iTRAQ subjected to LC-MS/MS. Both proteomes were combined, redundancies removed and confidently identified peptides with abundance changes of 2-fold or greater linked to protein identifiers. The lists of upregulated and downregulated proteins were submitted to the Database for Annotation, Visualization, and Integrated Discovery (DAVID) v6.7 and mapped to KEGG pathways. Significantly enriched major metabolic pathways were listed with proteins, soluble (Sol) or insoluble (Insol) fraction and fold change (FC) designated.

### DC restructure metabolism during IAV infection

Next, we sought to determine whether the changes in metabolic protein networks in our proteomic analysis were reflected in DC metabolism. To validate the protein network analysis and further characterize influenza-induced changes in metabolism, we quantified glucose, pyruvate, glutamine, and palmitic acid oxidation as previously described [14]. Isotopically labeled metabolic substrates were used to monitor generation of ^3^H_2_O from glycolysis or beta oxidation and ^14^CO_2_ from the pentose phosphate pathway, glutamine utilization, or TCA cycle. Glucose detritiation occurs early in glycolysis when fructose-6-phosphate is converted to fructose-1,6-bisphosphate. By monitoring glucose detritiation, we found that influenza infection significantly increased glycolysis (Fig 2A). PPP provides ribose 5-phosphate for nucleotide synthesis and NADPH as a cofactor for fatty acid synthesis. We quantified the oxidative portion of this pathway, following the release of CO_2_ from glucose-6-phosphate, and found the pentose phosphate pathway was unchanged by IAV infection (Fig 2B).

**Figure 2.**
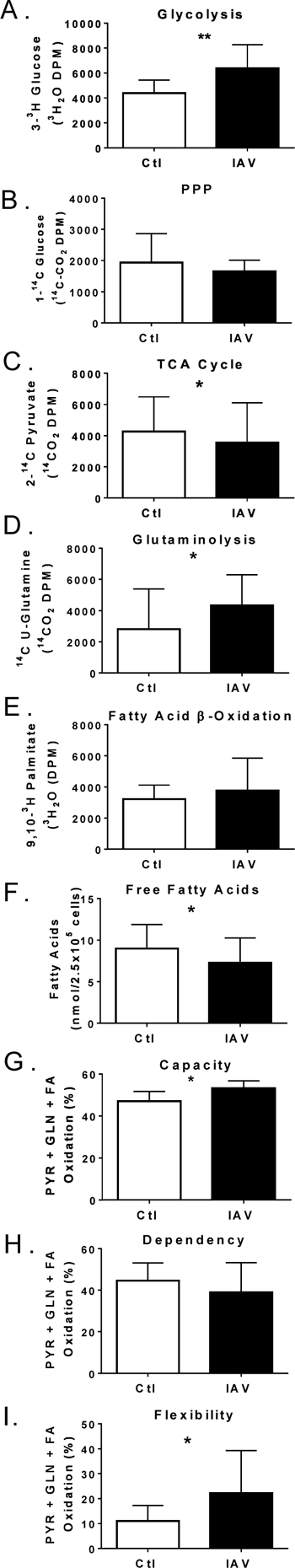
Metabolic changes in infected DC. DC were infected at MOI of 1 for 17 hours. (A-E) Use of isotopically labeled substrates, as indicated on the y axis, were used to monitor the generation of traceable products (^3^H_2_O or ^14^CO_2_) two hours after DC treatments substrates were quenched and allowed to accumulate overnight. Substrates were indicative of the following metabolic pathways: (A) [3-^3^H]glucose (glycolysis), (B) [1-^14^C]glucose (pentose phosphate pathway), (C) [2-^14^C]pyruvate (TCA cycle), (D) [U-^14^C]glutamine (glutaminolysis), and (E) [9,10-^3^H]palmitic acid (fatty acid oxidation). (F) DC was determined by a coupled enzyme assay releasing fluorometric product proportional to the fatty acids present and quantifiable with standards after treatments. (G-I) The rates of pyruvate, glutamine, or long chain fatty acids oxidation for respiration were calculated as the percentage of inhibition of oxygen consumption by UK5099, BPTES, or Etomoxir specific inhibitors of mitochondrial pyruvate carrier, glutaminase, and carnitine palmitoyl-transferase 1A. Capacity for a specific substrate to drive respiratory OCR was tested by determining baseline OCR then inhibiting the 2 off target substrates determining OCR then inhibiting import of the target metabolite. Percent capacity is one minus the baseline OCR less the off-target OCR divided by the baseline OCR less the OCR after all targets inhibited times 100. Dependency on a specific substrate was tested as above reversing the inhibitor sequence and the percent dependence was calculated by deducting the target OCR from the baseline and dividing by the baseline OCR less the OCR after all targets inhibited times 100. Fuel Flexibility was calculated as the difference between Capacity and Dependency. (G) The average capacity of uninfected or infected DC to use pyruvate, glutamine, or long chain fatty acids was determined. (H) The average dependence of uninfected or infected DC on the oxidation of pyruvate, glutamine, or long chain fatty acids was determined. I) The average flexibility of DC to alternate oxidation of pyruvate, glutamine, or long chain fatty acids was determined for uninfected or infected. The graphs represent the values of two (E&F), three (B-D), or four (A, G-I) independent experiments measured replicates) and presented as experimental mean +/- SD. Normality of these data were tested followed by the appropriate parametric (t-test) or nonparametric (Wilcoxon signed rank test) for normal distributions (A, B, E-H)) or not normal distributions (C,D,I), respectively. Asterisks correspond to p-values <0.05 (*) and a p-value < 0.01 (**).

Glycolysis is linked to the TCA cycle by the conversion of pyruvate to acetyl-CoA and CO_2_ by the pyruvate dehydrogenase complex. To assess TCA cycle, we quantified the aggregate release of ^14^CO_2_ from [2-^14^C]-pyruvate in 3 steps of the cycle. Consistent with the enrichment of this pathway in the decreased proteome, we found TCA cycle was significantly reduced following influenza infection of DC (Fig 2C). Based on the network and flux analyses, we further examined the pyruvate dehydrogenase complex. This is a large multifaceted protein complex. It was completely covered in the proteome, and most pyruvate dehydrogenase subunits decreased after infection (Table 3). In the canonical TCA cycle, in addition to pyruvate-derived acetyl-CoA, oxidation of fatty acids or glutamine can be used to produce ATP and intermediates. Glutamine is also a substrate for reductive carboxylation in the non-canonical reversal of the cycle. When we measured the release of isotopically labeled glutamine, we found DC increased glutaminolysis during IAV infection (Fig 2D). However, palmitic acid oxidation was unchanged in IAV infected DC concomitant to a decrease in intracellular free fatty acids (Fig 2E & F). Together, the increase in glycolysis and decrease in TCA cycle utilization of pyruvate is reminiscent of the Warburg effect that occurs in some cancers, in which the two major metabolic pathways become uncoupled to maintain ATP production while increasing metabolic plasticity [31–36].

**Table 3.** Proteomic coverage of the pyruvate dehydrogenase complex

| Subunit Function | Fold Change | Subunit Name | Fraction |
| --- | --- | --- | --- |
| E1 Regulatory (Activator) | -2 | Pyruvate dehydrogenase kinase 1 | Insol |
| E1 Regulatory (Activator) | -2 | Pyruvate dehydrogenase kinase 3 | Insol |
| E1 Catalytic | -1 | Pyruvate dehydrogenase beta | Sol |
| E1 Catalytic | -1 | Pyruvate dehydrogenase alpha | Sol |
| E1 Catalytic | -2 | Pyruvate dehydrogenase beta | Insol |
| E1 Catalytic | -2 | Pyruvate dehydrogenase alpha | Insol |
| E3 Non-catalytic | -2 | Pyruvate dehydrogenase component X | Insol |
| E2 Catalytic | 1 | Dihydrolipoamide S-acetyltransferase | Insol |
| E2 Catalytic | -2 | Dihydrolipoamide S-acetyltransferase | Sol |
| E3 Catalytic | 1 | Dihydrolipoamide dehydrogenase | Sol |
| E3 Catalytic | -2 | Dihydrolipoamide dehydrogenase | Insol |
| E1 Regulatory (Inhibitor) | -2 | Pyruvate dehydrogenase phosphatase | Insol |

The proteome was queried for components of the pyruvate dehydrogenase complex and subunits reported with corresponding functions, fold changes, names, and localization to soluble (Sol) or Insoluble (Insol). Summary of identified metabolic proteins that relocalized from soluble to insoluble fraction or vice versa. Fold enrichment significance was generated from DAVID functional annotation clustering and the biological significance indicated with p-value derived from KEGG pathways. Fold changes from non-unique proteins identified in the same metabolic pathway were averaged.

Some cancers and T cells display dynamic metabolism by acquiring more flexibility in metabolic substrate oxidation to facilitate macromolecule synthesis and ATP production [14, 15, 28, 37-42]. To determine the contribution of pyruvate, palmitic acid, and glutamate to DC bioenergetics, we monitored oxygen consumption via mitochondrial respiration using the XF Mito Fuel Flex Test as previously described [1, 43, 44]. This test was run on an XF Extracellular Flux Analyzer (Xfe96) that kinetically measures oxygen consumption rates (OCR) to monitor mitochondrial respiration over time while injecting stimulants or inhibitors. Xfe96 respirometry, in the context of inhibitor treatments, allowed us to probe DC energy utilization, substrate preference, and metabolic plasticity by selectively blocking mitochondrial oxidation of pyruvate, palmitic acid, or glutamate with UK5099, etomoxir, or BPTES, respectively. Note that while etomoxir can have off-target metabolic effects at high concentrations[45], it is a specific inhibitor of fatty acid oxidation in this context. Thus, the capacity of DC for oxidizing an isolated substrate was determined by quantifying the OCR after blocking the other two substrates, and, conversely, dependency on a single substrate was determined by inhibiting its oxidation (S2A-C Fig). We then combined the percent oxidation capacity or dependence and compared the infected and uninfected DC (Fig 2G & H). DC adopted a slight increase in capacity for oxidation of glucose, glutamine, and fatty acids to fuel mitochondrial respiration following infection (Fig 2G). However, IAV infection did not significantly alter DC dependence on mitochondrial oxidation of these substrates (Fig 2H). Together, capacity and dependency determine the flexibility of mitochondrial substrate oxidation (i.e. flexibility is the difference between the capacity to use a substrate and the dependency on that substrate). Overall, the mitochondrial flexibility for diverse metabolic substrates in DC was increased after IAV infection (Fig 2I).

Previously, we showed that neither the dependence nor capacity for oxidizing these mitochondrial substrates was dramatically impacted by IAV infection of epithelial cells, even though the trend for glutamine dependence increased [1]. However, when we calculated the individual contribution by substrate, we found quiescent DC had a significantly lower capacity and dependence on glutamine than on pyruvate and fatty acid oxidation (S2D Fig open bars). Yet, in response to IAV infection, DC significantly increased their capacity to use glutamate to fuel mitochondrial respiration (S2D Fig). When we blocked pyruvate entrance into the TCA cycle or fatty acid import for -Oxidation, we found a significant reduction in intracellular glutamine in IAV infected DC, presumably due to increased glutaminolysis in the absence of other oxidative substrates (S2E Fig). Next we wanted to determine if IAV infected DC would increase glutamate dehydrogenase enzyme activity to drive glutaminolysis in the absence of the other mitochondrial oxidative substrates. -ketoglutarate enzyme activity significantly increased when we blocked pyruvate with UK5099 in IAV infected DC (S2F Fig). Taken together, these data reveal that IAV-infected DC significantly increase glycolysis without increased shuttling of acetyl CoA from pyruvate into the TCA cycle, while also improving their capacity and flexibility for mitochondrial substrate oxidation, including markedly increasing glutaminolysis.

### IAV infected DC increase glycolytic bioenergetics similar to TLR stimulation

Substrate consumption for ATP production and product efflux vary with cellular metabolism and can be used to measure bioenergetic states. Thus, we also used the Xfe96 to determine the extracellular acidification rate (ECAR) as an indirect measure of glycolysis. Indeed, lactic acid excretion per unit time in glycolysis accounts for the majority of pH change in most cell types and glycolysis correlates well with extracellular lactic acid accumulation (i.e. R^2^ = 0.9101) [46, 47]. However, phagocytes employ radical generation and pH in signaling and innate effector functions, and therefore we used the glycolytic stress test to isolate glycolytic ECAR from lactate and non-glycolytic ECAR as previously described [47–49]. In keeping with our flux analysis, we found that influenza infected DC significantly increased glycolysis resulting in a mean difference of 26.5242 95% CI [1.62, 51.43] (Fig 3A). While previous reports indicated detritiation of D-[3-^3^H]- glucose underestimates glycolysis [50], we found the IAV-induced increases in glycolytic flux and bioenergetic measures of glycolysis were comparable (Figs 2A and 3A). We also infected DC with the 2009 pandemic H1N1 influenza A/California/04/2009 (CA04) and the H3N2 subtype influenza reassortment virus X31 that carries the HA and NA genes of A/Hong Kong/1/1968 in the background of PR8. Similar to the PR8 (Fig 2A-IAV), we found CA04 infected DC significantly increased glycolysis, resulting in a mean difference of 17.60 95% CI [6.017, 29.18] while the increase by X31 was not significant (S4A Fig).

**Figure 3.**
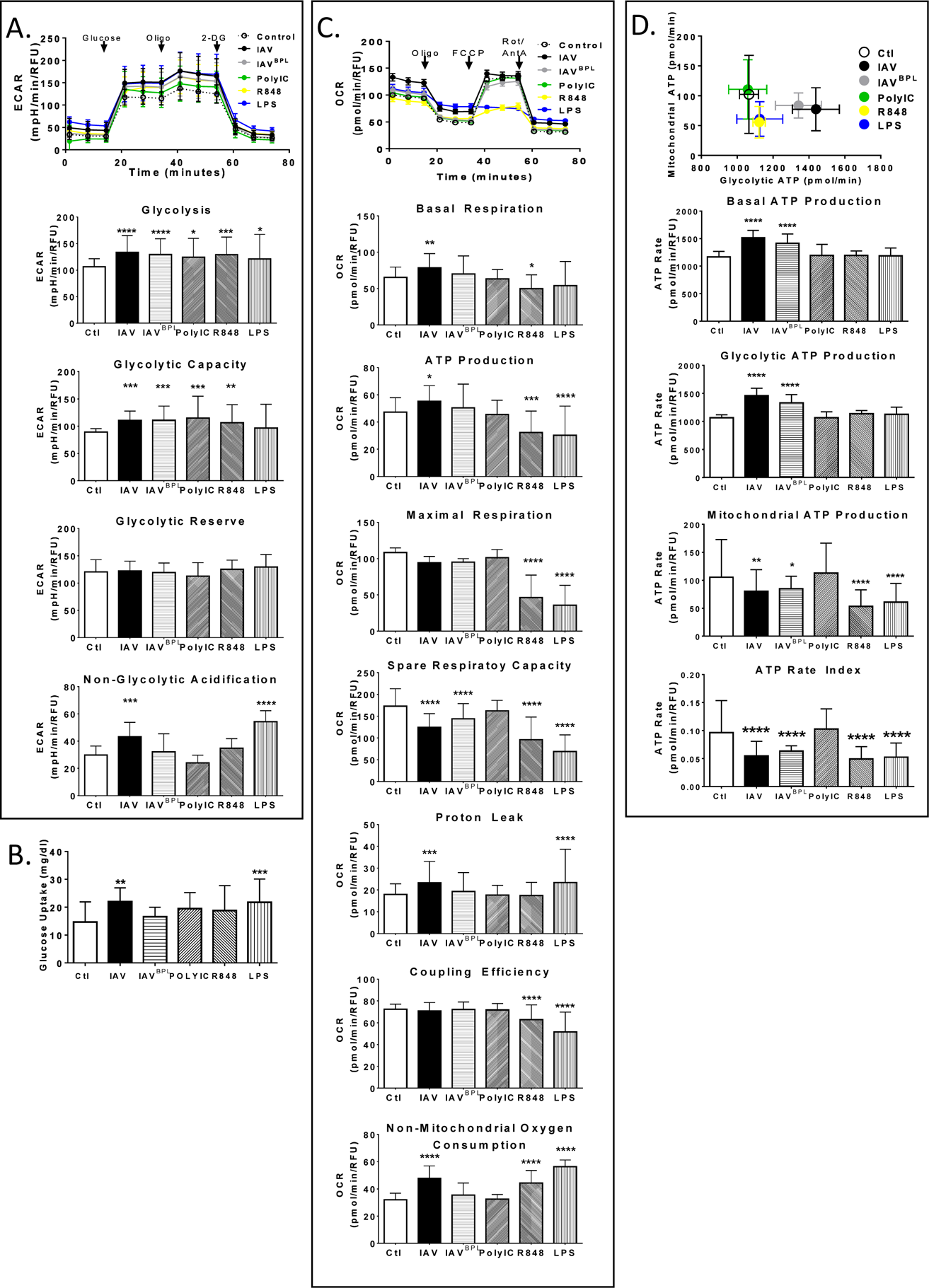
IAV infection induces distinct bioenergetics compared to TLR agonists primarily through aerobic glycolysis. (A-D) DC were infected or treated with TLA agonists lipopolysaccharide (LPS), polyinosinic polycytidylic acid (PolyIC), or Resiquimod (R848) for 17 hours followed by metabolic analysis with Seahorse Xfe96 Flux Analyzer. (A) Glycolytic function was tested with the Glycolysis Stress Test while monitoring real-time extracellular acidification rate (ECAR) with the Xfe96 metabolic analyzer during sequential injections of glucose, oligomycin (Oligo), and 2-Deoxy-D-glucose (2-DG) indicated by arrows. (B) Glucose uptake was monitored from the medium using a standard blood glucometer with glucose standard calibration curves. (C) Mitochondrial respiration was tested with the Mitochondrial Stress Test while monitoring oxygen consumption rates (OCR) in real-time with the Xfe96 metabolic analyzer during sequential injections of oligomycin (Oligo), carbonyl cyanide-p-trifluoromethoxyphenylhydrazone (FCCP), and a mixture of rotenone and antimycin A (Rot/AntA) indicated by arrows. (D) DC maximal mitochondrial ATP changes induced by oligomycin plotted against maximal ATP changes upon glucose depletion determined by respirometry using Xfe96. Each dataset represents the mean of 3 experiments +/- SD for bar graphs or +/- SEM for energetic traces. Statistical differences among means was found with ANOVA followed by Tukey and validated with Dunnett’s multiple comparison tests. Tukey test derived p values are symbolized by asterisks indicating adjusted p-, and **** p<0.0001).

We then sought to distinguish between cell-mediated responses to viral entry and innate recognition and to determine if DC restructure glycolysis equally with activating stimuli or tailor it in response to IAV infection. To this end, we treated DC with different TLR agonists and - propiolactone (BPL) inactivated IAV (IAV^BPL^) and compared these responses to active IAV. Influenza infects and replicates its genome producing viral proteins within DC, but virions are not released [21, 22]. In contrast, BPL treatment prevents genome replication but allows IAV to attach, fuse, and stimulate TLR7 in the endolysosome [51–53]. We used the TLR3 agonist polyinosinic polycytidylic acid (structurally similar to IAV dsRNA), TLR4 agonist lipopolysaccharide (LPS) to simulate DC detection of IAV-infected dead cells, and imidazoquinoline R848 (R848) for activation of both TLR7 and 8 and its adapter MyD88. Like IAV, these treatments significantly increased glycolysis (Fig 3A). ATP synthase inhibition with oligomycin decreases the ATP/ADP ratio thereby increasing glycolysis to maximal output, allowing us to calculate the glycolytic capacity of DC (i.e. maximal ECAR post oligomycin less non-glycolytic acidification). The glycolytic capacity was also increased by IAV infection, PolyIC, and R848, but not LPS (Fig 3A). The difference between the maximal glycolytic output and basal glycolysis is the glycolytic reserve, which remained constant across treatments (Fig 3A). The final injection of 2-DG allowed us to quantify any extracellular acidification occurring independent of glycolysis. We found that non-glycolytic acidification was significantly increased by active influenza infection and LPS treatment resulting in a mean difference from control of 13.16 and 23.29 respectively (Fig 3A and S3A Fig). We then quantified glucose uptake to determine if substrate import was coordinated with the increase in glycolysis. We found, in comparison to control, IAV and LPS significantly increased glucose uptake resulting in similar mean differences (Fig 3B and S3B Fig). Interestingly, glucose uptake in DC infected with active IAV was significantly higher than inactivate IAV^BPL^ (S3B Fig). Thus, in response to IAV infection DC increased both glucose uptake and glycolysis. However, in general these activators stimulated an uptick in glycolysis, which may provide additional energy and glucose-derived metabolic intermediates for macromolecule biosynthesis as both are necessary to fuel effector functions such as increased motility, phagocytosis, and cytokine production.

### IAV infected DC mitochondrial respiration is distinct from TLR stimulation

Next, we used respirometry with the Xfe96 to quantify and isolate individual components that contribute to mitochondrial respiration with the mitochondria stress test as previously described [54, 55]. This test is based on the principle that the amount of oxygen consumed during respiration is stoichiometrically related to the amount of ADP and substrate in respiratory oxidation. It probes respiration efficiency using sequential inhibition of complex V (ATP synthase) and complexes I and III of the electron transport chain with oligomycin and a mixture of rotenone and antimycin A, respectively, and the uncoupler FCCP to activate proton conductance. Using an analysis of variance (ANOVA) on all the treatments yielded significant variation among conditions (p < .0001) and with a post hoc Tukey’s Honest Significant Difference test we showed that control untreated and IAV infected DC differed significantly at p < .0001 (Fig 3C). Only IAV increased respiration, resulting in a mean difference of 22.51 pmol/min/RFU with a 95% CI [3.48, 41.54] between IAV and control (S3C Fig). Conversely, PolyIC, LPS, and R848, all reduced basal respiration by mean differences of 2.3, 7.6, and 12.4 pmol/min/RFU, respectively, but only the R848 group difference was significant (Fig 3C). Accordingly, we also showed IAV differed significantly from PolyIC, R848, and LPS (S3C Fig). Similarly, we found modest but reliable increases in respiration when DC were infected with either the CA04 or X31 that are H1N1 and the H3N2 subtypes, respectively (S4B Fig). Therefore, in response to IAV infection, DC significantly increase basal respiration and this response is significantly different from PolyIC, LPS, or R848 treatment.

After three basal respiratory measures were taken the ATP synthase inhibitor oligomycin was added to isolate OCR linked to ATP production. IAV infection induced a modest increase in oxygen consumption linked to ATP synthase activity in DC, whereas LPS and R848 treatments significantly reduced respiratory ATP production (Fig 3C). Like basal respiration, these opposing responses resulted in significant differences in respiratory ATP production between the IAV infected and PolyIC, LPS, or R848 treated DC (S3C Fig). To maximize respiratory OCR, we then uncoupled oxygen consumption from ATP production with the addition of the protonophore FCCP. This allowed us to calculate the maximal oxygen consumption of DC for respiration, which was significantly impaired by LPS or R848 treatment but unperturbed by IAV infection (Fig 3C). These three measures demonstrate that IAV infected DC increase basal oxygen consumption that is associated with mitochondrial respiration whereas in response to R848 and LPS treatments both ATP production and maximal respiration are impaired.

The difference between maximal and basal respiration, or the spare respiratory capacity, indicates how closely cells are operating near their OXPHOS threshold. Untreated and PolyIC treated DC displayed similarly low basal and high maximal respiration, resulting in similar spare respiratory capacity (Fig 3C). In contrast, DC treated with either R848 or LPS failed to respond to FCCP resulting in very low maximal respiration and spare respiratory capacity (Fig 3C and S3C Fig). By comparison, IAV infected DC increased OCR when FCCP activated proton conductance but had no change in maximal respiration (Fig 3C top). This was due to the significantly higher basal rate, near DC maximal respiration, thereby reducing spare respiratory capacity of IAV infected DC. Indeed, the mean difference in IAV infected DC and untreated controls spare respiratory capacity is −40.62 with 95% CI [-69.16, 12.09], a significant difference P < 0.0001 (Fig 3C and S3C Fig). Therefore, IAV, LPS, and R848 treated DC had significantly reduced spare respiratory capacity, but different respiratory responses. Thus, due to IAV inducing an uptick in basal respiration the infected DC were already operating closer to their maximal respiratory output thereby diminishing their capacity to increase respiration, whereas LPS stimulated DC had a well-documented respiratory collapse and did not respond to uncoupling [18, 19].

The final inhibitor combination of rotenone and antimycin A blocks proton transport in the electron transport chain by inhibiting complex I and III for the quantification of proton leak and residual OCR from other sources (e.g. NADPH oxidases, arachidonic acid metabolism, NOX enzymes, etc.). OXPHOS proton leak is the respiratory OCR that is not coupled to respiratory ATP production and, under these conditions, it is unresponsive to fluctuations in substrate oxidation and unaffected by variations in ATP turnover [56]. IAV infection and LPS treatment showed significantly more proton leak than controls (Fig 3C). Mitochondrial coupling efficiency is the amount of ATP produced through the electron transport chain per reduction of consumed oxygen, it is similar to the traditional phosphate oxygen ratio and both are sensitive to mitochondrial dysfunction [56]. Proton leak does respond to respiratory dysfunction from uncoupling and can indicate severe mitochondrial damage [56]. However, we found that the respiratory coupling efficiency of DC was stable for IAV, IAV^BPL^, and PolyIC as opposed to LPS and R848 treatments that were significantly reduced (Fig 3C). Taken together, the impaired FCCP response of both LPS and R848 and diminished coupling efficiency indicate mitochondrial damage, which is in keeping with current reports on the impact of LPS on DC respiration [18, 19].

We also found that the non-mitochondrial respiration oxygen consumption rates were significantly increased by active IAV influenza infection, R848, and LPS treatment (Fig 3C). This was not surprising, given that activated phagocytes produce reactive oxygen species (ROS) from inflammatory enzymes cyclooxygenases, lipooxygenases and NADPH oxidases, NADPH oxidases alone can account for up to 90% of total cellular oxygen consumption in these cells [57]. ROS damages mitochondria and varies with non-mitochondrial respiration OCR [58]. Thus it is possible that R848 and LPS over activate proinflammatory enzymes that divert oxygen away from respiration creating a significant increase in non-mitochondrial OCR concomitant to reducing OCR in basal respiration lowering ATP production and potentially damaging mitochondria as is evident by the lack of FCCP response creating significant drops in maximal and spare respiration along with decreased coupling efficiency, as well as proton leak for LPS (Fig 3C). Non-mitochondrial OCR rose 175% in DC with LPS treatment and was significantly higher than all other treatment groups (Fig 3C and S3C Fig). Combined, these measures provide an indication of the bioenergetic state of activated DC. Impairment of the electron transport chain will lower coupling efficiency, mitochondrial ATP production, and spare respiratory capacity while increasing proton leak. However, cells can still meet their metabolic demands with proton leak and in the presence of heightened basal respiration that lessens the dynamic range and prevents dramatic bursts in respiration [58].

In lieu of mitochondrial impairment, these measures of pyruvate flux, glutamine flux, and respirometry showed that influenza infection significantly reduced normal pyruvate oxidation while increasing glutamine oxidation for fueling the TCA cycle. This makes sense given 17 hours after IAV infection DC had significantly depleted glucose in their surrounds (i.e. a mean difference of −6.680 mg/dl glucose with 95% CI [-12.24, −1.115]) (Fig 3B). Further, when glucoses derived pyruvate is lacking DC are known to shift to glutaminolysis to fuel TCA cycle to support OXPHOS [19, 58-62]. Thus, even though we find IAV infection results in significant increases in DC glycolysis it appears to uncouple from the TCA cycle, via reduced pyruvate dehydrogenase, concomitant to a modest increase in glutaminolysis acting to fuel OXPHOS related oxygen consumption without evidence of mitochondrial damage as measured by FCCP response and coupling efficiency.

### IAV infection DC increase ATP production via glycolysis distinct from TLR agonists

Normally, the majority of cellular ATP is derived from OXPHOS downstream of glycolysis and the TCA cycle. However, glucose import, oxidation, and basal glycolysis significantly increased after infection of DC, concomitant to declining pyruvate oxidation while respiration and respiratory capacity and flexibility increased (Figs 2 and 3). Collectively these findings support a transition of DC metabolism to more aerobic glycolysis and non-canonical TCA, reducing the efficiency of ATP production per oxidized substrate. To determine the source of ATP production in IAV infected DC, we measured total ATP production rates and that derived from glycolysis or mitochondrial OXPHOS. Changes in the relative output of ATP from glycolysis and mitochondrial respiration often reflects the cell’s metabolic phenotype. Consistent with our previous bioenergetic measures of glycolysis and respiration, we found that DC paired into three distinct metabolic phenotypes: hyper metabolic IAV and IAV^BPL^ (black and gray), metabolically impaired R848 and LPS (yellow and blue), and normal Ctl and PolyIC (open circle and green) (Fig 3D top). DC significantly increased total ATP production in response to IAV infection, irrespective of BPL inactivation, (Fig 3D). Indeed, following IAV infection DC had a striking 39% increase in glycolytic ATP production resulting in a mean difference of 418.6 pmol/min/RFU with 95% CI [297.1, 540.1] when compared to controls (Fig 3D & S3D Fig). In contrast, IAV infection diminished mitochondrial ATP while R848 or LPS treatment significantly reduced it (Fig 3D & S3D Fig). One hallmark indicator of a fundamental shift in metabolic phenotype is a change in the ATP ratio from mitochondrial to glycolytic ATP production. Due to the dramatic and very significant increase in glycolytic ATP production concomitant to modest changes in mitochondrial ATP production following infection, the ATP rate index significantly decreased DC following infection (Fig 3D bottom). This indicates DC change to a more glycolytic and less oxidative phenotype following IAV infection. In contrast, DC activated with either LPS or R848 also exhibit a significant decrease in the ATP rate index, but this was due to a very significant reduction in mitochondrial ATP production without a change in glycolytic ATP (Fig 3A & D). Collectively these metabolic and bioenergetic studies reveal that DC re-wire their metabolism in response to IAV infection in a manner that resembles Warburg metabolism in cancer and is distinct from TLR4 and TLR7/8 agonist treatment.

### Transcriptional regulation of DC metabolic phenotype

One transcription factor promoting the highly glycolytic and glutaminolytic metabolic transition in cancer and activate T lymphocytes is c-MYC [14, 63-67]. DC express variable levels of three Myc family members (c-MYC, n-MYC, and l-MYC) that regulate immune cell development, differentiation, and activation [14, 68, 69]. We found IAV infection induced transient c-MYC expression peaking at 4 hours (Fig 4A). This increase in c-MYC expression in IAV-infected DC was reflected in increased protein levels by 17 hours (Fig 4A inset). In many cancers, some c-Myc driven, this metabolic phenotype is also associated with sensitivity to glucose and glutamine deprivation that results in rapid cell death if these metabolites are restricted and is often referred to as glucose and glutamine addiction [63, 65-67, 70-72]. We previously found c-Myc transiently increased in IAV infected epithelial cells and glucose and glutamine restriction caused an approximate 50% decrease in their viability [1]. Therefore, we quantified DC viability and cell death using CalcienAM and ethidium bromide homodimer (EthD-1) per total cell content via DAPI. Cell death was minimal, impacting less than 10% of DC, irrespective of metabolite restriction or treatment (S4C Fig). In the absence of glucose, the proportion of viable DC dropped (S4C Fig open bars). However, neither infection nor activation with TLR agonists increased cell death (i.e. reduce the live to dead ratio) in the absence of glucose (S4C Fig). The same was true when glutamine was restricted apart from R848 and LPS, these treatments significantly reduced cell death thereby increasing the live to dead ratio (S4C Fig). In contrast to epithelial cells, removal of both glutamine and glucose had no adverse effect on IAV-infected DC (S4C Fig open and solid bars). LPS was the only treatment to slightly increase DC cell death, from 6.4% or 7.1% for the complete or deplete controls, to 9.8% for DC treated with LPS in the absence of both glutamine and glucose (S4C Fig red vertical stripes). We observed high levels of viability with very little cell death in all treatments, indicating that metabolic differences were not simply due to population decline or adverse effects from treatment with high concentrations of agonists. Further, the viability of infected DC was more stable with varying metabolite levels than we previously observed in IAV-infected respiratory epithelial cells [1]. These findings were surprising, given the transient in c-Myc (Fig4A), but were in keeping with DC not developing substrate dependency (Fig 2H) and likely due to the significant increase in metabolic flexibility and capacity to utilize diverse substrates that DC acquire with infection (Fig 2G & I).

**Figure 4.**
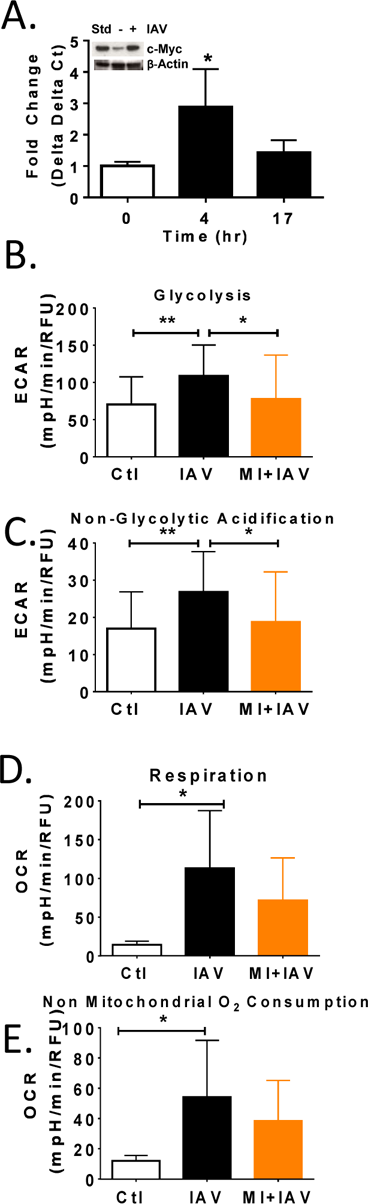
c-Myc inhibition blocks IAV induced glycolysis. (A) Up to 17 hours PI qPCR and immunoblotting of RNA and lysates were performed and quantification of target gene expression method. One representative blot is presented in inset. (B-E) DCs were seeded in Seahorse XFe-96 plates and pretreated with cMyc inhibitor (MI) for 4 hours then IAV infected for 17 hours. (B-C) Glycolytic function was tested with the Glycolysis Stress Test while monitoring real-time extracellular acidification rate (ECAR) with the Xfe96 metabolic analyzer during sequential injections of glucose, oligomycin (Oligo), and 2-Deoxy-D- glucose (2-DG). Data represent means ± SD of 4 experiments. (D-E) Mitochondrial respiration was tested with the Mitochondrial Stress Test while monitoring oxygen consumption rates (OCR) in real-time with the Xfe96 metabolic analyzer during sequential injections of oligomycin (Oligo), carbonyl cyanide-p-trifluoromethoxyphenylhydrazone (FCCP), and a mixture of rotenone and antimycin A (Rot/AntA). These graphs represent the mean values of 3-4 independent experiments +/- SD. Statistical differences among means wese found with ANOVA followed by Tukey’s multiple comparisons test with asterisks indicating adjusted p- and **** p<0.0001).

To determine if DC require c-MYC activity to modulate their metabolic response to infection, we pretreated DC with a c-MYC inhibitor (MI) also known as 10058-F4, for 4 hours, then infected them and measured glycolysis and non-glycolytic acidification. After infection DC increased glycolysis by 54.3%, but c-MYC inhibition caused this to drop significantly to 10.2% similar to uninfected levels (Fig 4B). MI treatment also reduced non-glycolytic acidification following IAV infection (Fig 4C). Increases in c-MYC primarily boost metabolism through the glycolytic pathway by increasing levels of lactate dehydrogenase A, glucose transporter 1, phosphoglucose isomerase, phosphofructokinase, glyceraldehyde-3-phosphate dehydrogenase, phosphoglycerate kinase, and enolase [73–75]. We found many of these proteins increased in the proteome, including-enolase (S1 A&D Fig). However more recent studies have implicated c-MYC in regulating mitochondrial metabolism and biogenesis as well [67, 76-78]. To determine if c-MYC was regulating the modest IAV induced changes in respiration we quantified basal respiration and non-mitochondrial oxygen consumption and found MI treatment had less effect (Fig 4D-E). Thus, in response to IAV infection, DC increased c-MYC expression and glycolysis and inhibiting c-MYC activity blocked the IAV induced glycolytic increase.

### Metabolic rewiring in IAV infection is necessary for DC effector functions

DC are controllers of innate and adaptive immunity and they are the primary immune cell population responsible for inducing the IAV virus specific CD8 T cell response requisite for viral elimination. Thus DC migration from the lungs to the lymph nodes is a critical first step in bridging the innate and adaptive response to IAV [5]. Increases in DC trafficking occur quickly after IAV infection and peak in vivo around 18 hours [79, 80]. We used a fluorescent cell migration assay to quantify DC motility in response to IAV infection. We found a very significant increase in DC migration after IAV infection (Fig 5A black). Myc inhibition had no effect on the migration of uninfected DC, while completely ablating IAV-infected DC migration (Fig 5A orange). Similarly, inhibiting the oxidation of pyruvate via the TCA cycle with UK5099 had no impact on the migration of uninfected DC (open bars), but significantly reduced infected DC migration by approximately 30% (Fig 5A red). Nonetheless, this migration was significantly higher than the controls (Fig 5A). Inhibiting fatty acid or glutamine oxidation in the mitochondria resulted in the same changes in DC migration as inhibiting pyruvate, primarily a significant decline in infected DC mobility that remained much higher than uninfected controls (Fig 5A blue and green). We then sought to determine if phagocytic activity of infected DC could be modulated by targeting these same pathways and found no effect (S5B Fig). While the role for DC phagocytosis post infection is unclear, and likely minimal we also found no differences among uninfected drug treated controls (S5B Fig open bars). Thus, inhibiting c-MYC ablates IAV-induced DC migration while blocking mitochondrial oxidation of pyruvate, glutamine, or fatty acids, significantly impairs it.

**Figure 5.**
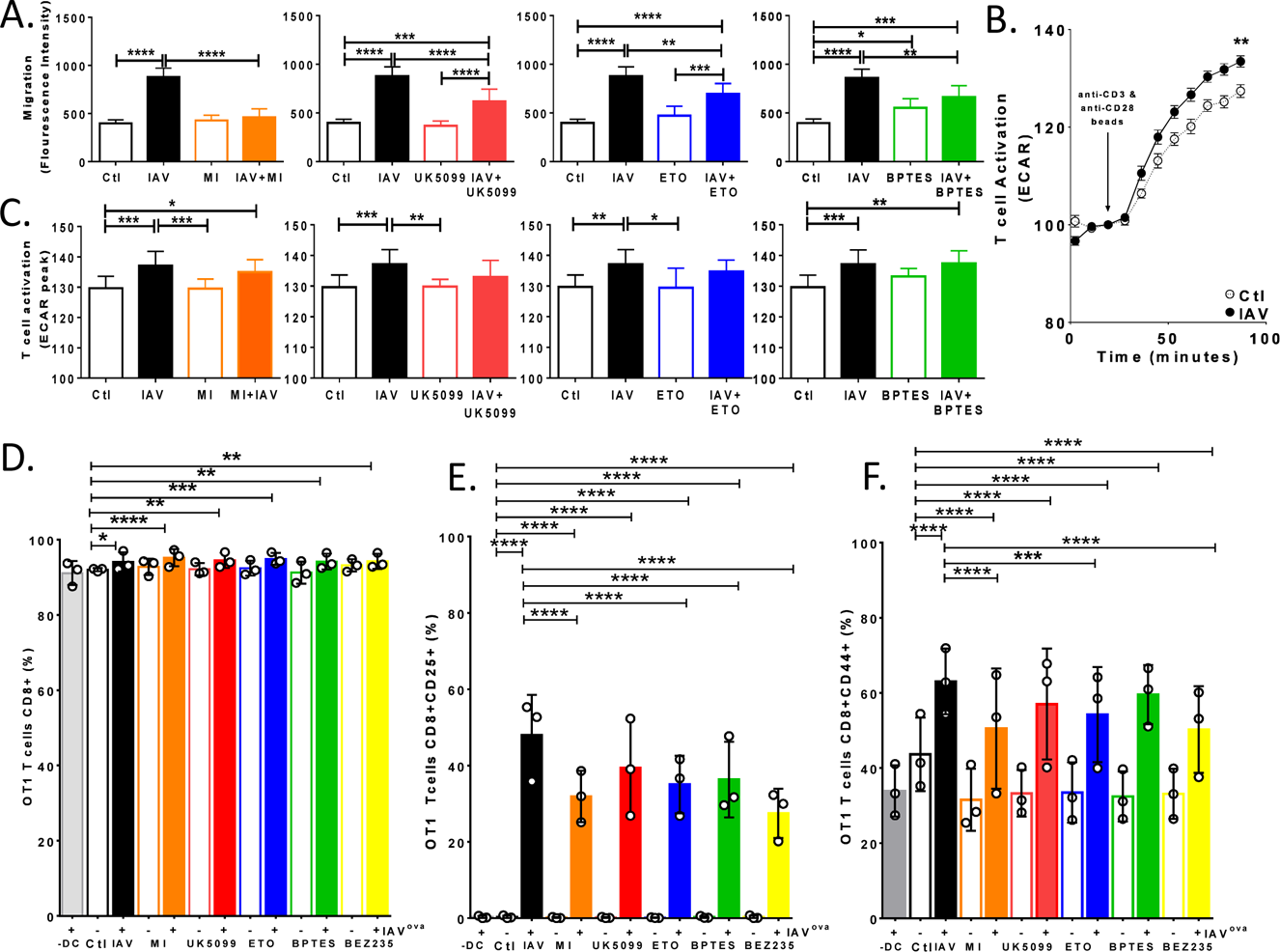
Limiting glycolysis or mitochondrial oxidation of pyruvate, glutamine or long chain fatty acids impairs infected DC function. (A) DC were seeded on precoated Oris™ 96- well migration plate allowed to adhere 18+ hours followed by 4 hours of inhibitor treatment (2uM c-Myc inhibitor, 3uM BPTES, 4uM etomoxir or 2uM UK5099) plug removal and infection (MOI = 5) for 17 hours. Cells were stained with calcein AM (2µM), viability and migration were determined with fluorescence at 485/528nm). The graph represents the mean values of 3 experiments +/- SD. (B-C) CD8+ T cells were isolated by negative depletion from fresh splenocytes of female C57BL/6 mice and co-cultured with DC at 5:1 ratio T cells to DC for 24 hours. Basal ECAR was established and T cell activation by anti-CD3/CD28-coated DynaBeads was monitored for 2 hours and maximal ECAR determined. (B) One representative graph is shown from 3 independent experiments. (C) The graph represents the mean values of 3 experiments +/- SD. (D) CD8+ T cells were isolated by negative depletion from fresh splenocytes derived from C57BL/6- Tg(TcraTcrb)1100Mjb/J (OT1) and co-cultured with DC at 5:1 ratio of T cells to DC for 24 hours. T cell were then stained for CD8, CD45.1, CD25, and CD44 and enumerated with flow cytometry. We selected CD8 and CD45.1 positive and then gated on CD25 and CD44. The graph represent the mean values of 2 experiments +/- SD. Statistical differences among means was found with ANOVA followed by Tukey’s multiple comparisons test with asterisks indicating adjusted p-values (* p ≤ 0.05, **p ≤ 0.001, and **** p < 0.0001).

Initial studies of DC function in vitro revealed that IAV can infect and stimulate DC maturation into potent antigen presenting cells that elicit robust IAV-specific T cell response relative to other APC without the addition of exogenous cytokines [81–83]. To delineate the roles of c-MYC, glycolysis, and mitochondrial substrate oxidation in DC maturation, we inhibited each and then infected the DC with IAV for 17 hours. The DC were then transferred to plates of freshly purified, wild type CD8 and CD4 T cells from naïve or homologously primed C57BL/6 mice at a ratio of 1 to 5 and co-cultured for 24 or 48 hours. It is well documented that T cells rapidly respond to activation with an immediate and robust glycolytic switch that increases ECAR and can be monitored to determine their activation state and distinguish T cell responses after anti-CD3/CD28 beads stimulation [14, 84-88]. Indeed, using this T cell activation assay as previously described [85–88], we found primary splenic T cells rapidly responded to anti-CD3/CD28 bead stimulation with clear and measurable increases in ECAR (Fig 5B). Furthermore, this ECAR peak was significantly increased when these T cells were co-cultured with IAV-infected DC as compared to uninfected controls (Fig 5B). Irrespective of inhibition of glycolysis or glutaminolysis with prior treatment of DC with c-MYC inhibitor or BPTES respectively, IAV infected DC effectively primed T cells resulting in a significant increase in T cell activation with bead stimulation (Fig 5C orange and green). In contrast, inhibition of pyruvate or fatty acid oxidation prevented effective IAV induced DC priming of T cells resulting in activation peaks that were the same as uninfected controls (Fig 5 C red and blue).

Following DC-T cell co-culture, we assessed CD8+ T cell populations staining positive for CD8, CD25 and CD44 with flow cytometry (S5C Fig). Only etomoxir treatment reduced CD8+ T cells (S5D Fig). DC that were pretreated with metabolic inhibitors reduced the CD25+ T cell population following co-culture (S5E Fig). However, due to the response heterogeneity of the T cell population from naive mice we move to a transgenic system. We purified CD8+ T cells from spleens of C57BL/6-Tg (TcraTcrb)1100Mjb/J (OT1) mice that contain transgenic inserts for Tcra- V2 and Tcrb-V5 genes. OT1 T cell receptor interacts with the ovalbumin amino acid sequence SIINFEKL, corresponding to residues 257-264, in the context of H2Kb. Dendritic cells were then vehicle or drug treated and infected for 17 hours with a PR8 influenza virus reverse engineered to express the SIINFEKL OVA peptide in the neuraminidase stalk (IAV^OVA^) that was generously provided by Dr Richard Webby as previously described [89]. We used three controls: T cells without DC (-DC), T cells co-cultured with uninfected and untreated control DC (Ctl), and T cells co-cultured with untreated DC infected with IAV^OVA^. We found DC infection with IAV^OVA^significantly increased the population of CD8+ T cell irrespective of drug treatment (Fig 5D and S5E Fig). We used cell surface staining and flow cytometry to quantify to CD25+ and CD44+ populations of CD8 T cells. In keeping with our previous results, we found that modulation of DC metabolism can significantly reduce the population of CD8+ T cells expressing the early activation marker CD25 compared to IAV^OVA^ alone (Fig 5E and S5F Fig). However, in the transgenic OT1 system IAV^OVA^ infection induced a significant rise in the proportion of T cells expressing CD25 irrespective of drug treatment (Fig 5E and S5F Fig). Similarly, metabolic inhibition of DC was not strong enough to abolish the significant increase in expression of CD44 on OT1 T cells in response to co-culture with IAV^OVA^ infected DC (Fig 5F). Importantly, in each of the 3 separate experiments DC treated with Myc inhibitor, etomoxir or BEZ235 significantly lower the population of CD44+ relative to IAV (Fig 5F and S5D&E). Based on our previous studies in IAV infected epithelial cells and DC, we knew the PI3K/mTOR inhibitor BEZ235 would block the IAV induced increase in glycolysis (1). Thus, we wanted to compare T cell response to DC treated with these inhibitors and found that BEZ235 significantly reduced the CD25 or CD8 positive T cells similar to Myc inhibitor (Fig 5E&F orange and yellow). Collectively these data show specific metabolic pathways are critical to DC functions such as motility and T cell activation. Namely, limiting glycolysis ablated IAV induced DC migration, a critical step for DC to get to the lymph nodes to initiate the adaptive response, and reduces CD25/CD44 positive CD8 T cell populations. In contrast, access to diverse fuels for mitochondrial respiration appears to have less effect on migration but play an more important role in rapid T cell activation and induction of the CD25 positive CD8 population.

### Lung TipDC globally reprogram metabolism in response to IAV infection in vivo

We have speculated that glycolysis and specific mitochondrial substrates that fuel OXPHOS are critical for DC migration and T cell priming in situ based on our in vitro models, and asserted the former is particular likely to affect immune response during influenza infection given DC translocation is a major determinant of outcome [79, 80, 90]. To determine if these metabolic pathways were relevant in vivo we intranasally infected C57BL/6 mice with IAV or mock for 9 days and removed the lungs. We isolated immune cells from the lungs and sorted DC based on CD11b^hi^, Ly6c/GR-1^hi^, MHCII ^hi^ DC surface marker staining (TipDC). TipDC are known for being inflammatory and producing TNF- and inducible nitric oxide synthase. In previous studies we showed this DC subset presents antigen to CD8 T cells in the lungs of mice infected with influenza A viruses and found TipDC are required to produce adequate influenza-specific CD8+ T cell proliferation in the infected lung to mediate viral clearance [91]. We purified RNA from the TipDC, generated cDNA libraries and subjected them to sequencing using the HiSeq2000 system (Illumina, San Diego CA) as previously described [92]. The paired end RNA-Seq data was aligned, concordant pairs were kept (MAPQ>10), a read number assigned to each gene (45,142 reads) and all genes with less than 3 counts or FDR > 0.05 were excluded. Thus, after redundancies were aggregated by mean, there were 13,285 confidently identified transcripts (Table A in S2 File). To evaluate the group trends, sample uniformity and identify potential outliers unsupervised multivariant principal component analysis (PCA) was employed. F1 and F2 explained the total variance with a cumulative percent variability of 91.35% (S6A Fig). We find the IAV has similar component 1 values and the controls have similar component 1 values, indicating the samples within each group have similar gene expression levels (S6A Fig). The double filtered volcano plot in Supplemental Figure 6B describes the selection criterion with the statistical effects on the y-axis and biological effects on the x-axis (dotted lines mark fold change 2.7 on the x axis and p- on the y axis).

These data were then independently k-means clustered followed by ascendant hierarchical clustering based on Euclidian distances. Genes with very little variability were non-specifically filtered using a standard deviation threshold of 50% to simplify the graph. The data matrix was rearranged according to the corresponding clustering with similarity proportional to a closer spatial relationship for DC columns and transcript rows. These clustering results were also represented via a dendrogram displayed vertically for genes and horizontally for TipDC. This resulted in clean separation of the TipDC from day 0 and the TipDC from day 9 of the IAV infection (Fig 6A). The data values of the permuted matrix were replaced by corresponding color intensities based on interquartile range with color scale of blue to red through white resulting in a heat map that vividly demonstrates the difference in gene expression of these infected and uninfected TipDC (Fig 6A).

**Figure 6.**
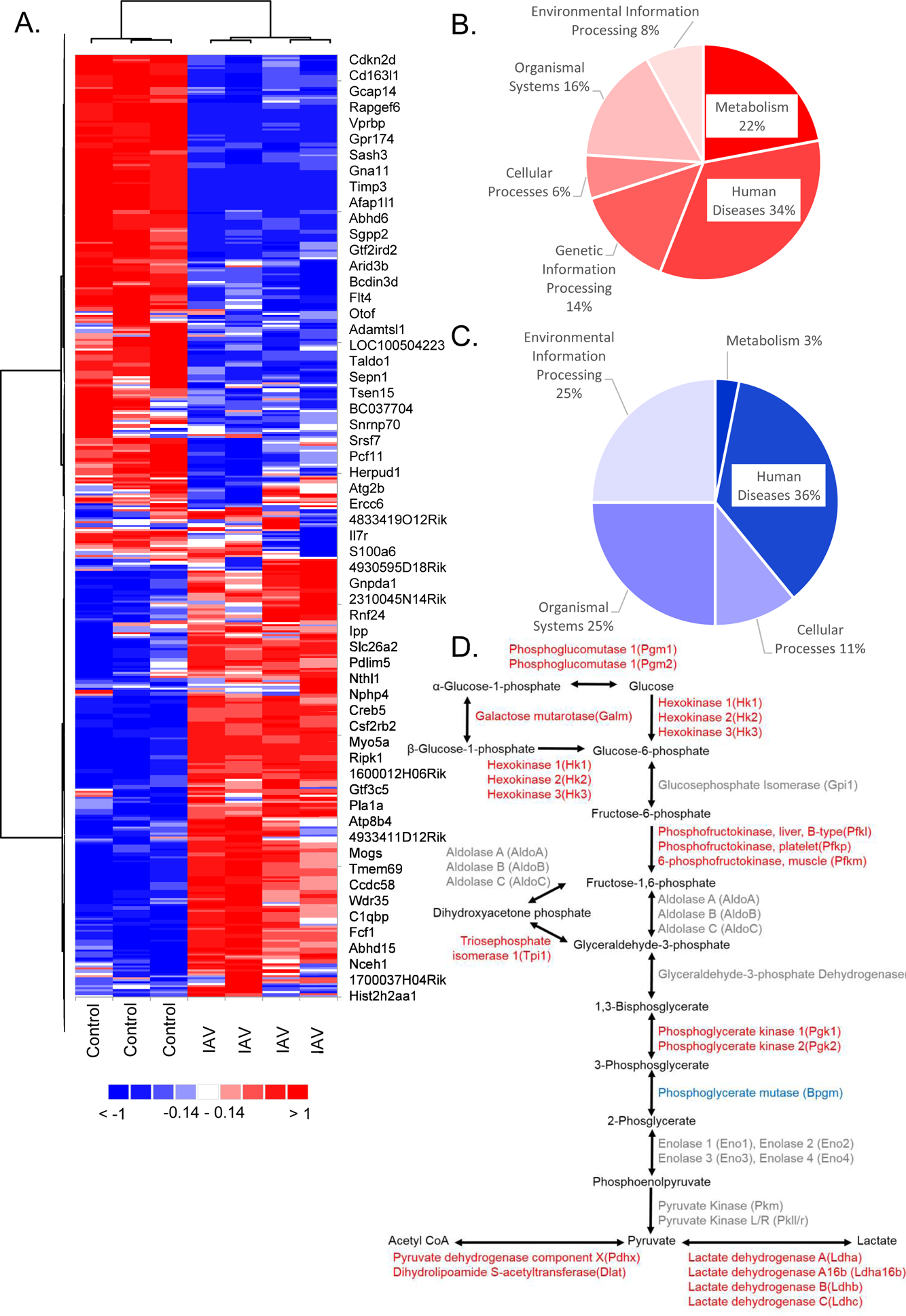
TipDC reprogram metabolism after accumulating in the lungs of IAV infected mice. At zero or nine days following intranasal infection of mice with IAV (PR8) lungs were homogenized and cells extracted. Cells were antibody stained for TipDC surface markers CD11b, Ly6c, GR-1, and MHCII. TipDC controls (Ctl) from day 0 were sorted based on high levels of CD11b, Ly6c, and GR-1. TipDC from day 9 of the IAV infection (IAV) were sorted based on high levels of CD11b, Ly6c, GR-1, and MHCII. cDNA libraries were generated from RNA and sequenced. (A) Confidentially identified transcripts were k-means clustered followed by ascendant hierarchical clustering and genes with standard deviation threshold < 50% were removed. The data values were replaced by corresponding color intensities of blue to red through white based on interquartile range. (B-D) Significance in gene expression differences were determined using Tukey honest significant difference test for multiple comparisons with a Benjamini-Hochberg post-hoc false discover rate correction. TipDC genes that were either upregulated (red) or downregulated (blue) following IAV infection were mapped to KEGG pathways and significantly enriched pathways identified and sorted by KEGG Class (B-C). All isoforms of the central enzymes in glycolysis from the TipDC transcriptome are presented and colored including those that were unchanged (gray).

We then used the OMICS data analytics package (Addinsoft, New York) to identify gene transcripts that were significantly affected by IAV infection in the context of specific criteria. We used the Differential Expression Analysis tool to apply a non-specific filtering threshold of 50% based on standard deviation and identifying significant differences in expression based on p-values generated from a parametric comparison of differences (Tukey honest significant difference test for multiple comparisons) with a post-hoc false discover rate correction (Benjamini-Hochberg). When compared to day 0 uninfected controls (Ctl), we found 4,788 transcripts were significantly altered on day 9 of IAV infection (Table B in S2 File). Gene identifiers of transcripts that increased with infection were input to DAVID for KEGG pathway enrichment analysis. Fifty pathways were significantly enriched, many of these were expected for TipDC in the IAV infected lung such as those governing immune response (e.g. Lysosome, Toll-like receptor signaling pathway, phagosome, TNF signaling pathway, endocytosis, Fc gamma R-mediated phagocytosis, etc.) as well as Influenza A (Table C in S File). Of the 50, global Metabolic Pathways (mmu01100) was the second most significantly enriched (Table C in S2 File). Like the in vitro differentiated DC proteome, we found pathways within the Metabolism KEGG Class comprise a large proportion of the increased TipDC transcriptome (Fig 6B). However, for the TipDC the Human Disease class was larger (Fig 6B). This is not surprising, given it includes infectious disease and cancer accounting for 8 and 6 out of 17 pathways, respectively. We then input the genes from the 11 significantly enriched metabolic pathways into DAVID to identify the diversity of the metabolic pathways associated with these transcripts that were significantly increased following IAV infection (Table D in S2 File). Glycolysis, oxidative phosphorylation, purine metabolism, and amino acid metabolism are some of the pathways these proteins regulate. Figure 6D depicts the enzymes and products of glycolysis with the transcripts that were significantly altered by IAV infection, we see expression of the majority of these key enzymes was upregulated. Thus, among TipDC transcripts that significantly increased nine days following IAV infection we find eleven metabolic pathways were significantly enriched, including the global Metabolic pathway (p < 0.00001) and glycolytic enzymes were significantly enriched (p < 0.00001).

In contrast, metabolism only made up 3% of the KEGG Classes and the global Metabolic pathways (KEGG ID mmu01100) was not found in the enrichment analysis of the TipDC transcripts that decreased following IAV infection (Fig 6C and Table E in S2 File). However, we did find two pathways associated with metabolism in the decreased transcriptome (Purine metabolism and Metabolism of xenobiotics by cytochrome P450). Many pathways enriched in the down regulated transcriptome reflect pathways one would anticipate being suppressed in TipDC on day 9 of the infection (e.g. leukocyte transendothelial migration, extra cellular matrix (ECM)- receptor interactions, cell adhesion molecules (CAMs), focal adhesions, etc.). We were surprised to see Influenza A pathway enriched in the transcripts that decreased with IAV. On closer analysis we found it was differentially regulated, as was Cytokine-cytokine receptor interaction, and Fc gamma R-mediated phagocytosis (Table F in S2 File). These include increased expression of viral sensors TLR4, TLR3, IFNAR1, IFNAR2 and related signaling molecules such as MyD88, STAT1, and STAT2 concomitant to decreased expression of FAS, FASLG and TRAIL along with NS1 regulated proteins NFX1, NFX 2, NFX 3, NFX5, PIK3CA, PIK3CB, PIK3CD, PIK3R1, PIK3R2, PIK3R, and PIK3R3. While the glycolysis pathway was significantly enriched in the increased transcripts, when we input all transcripts that were significantly altered by IAV infection we found a few proteins related to glycolysis were decrease (Fig 6D blue).

Importantly, when we compare the in vitro and in vivo IAV infected DC datasets we find many similarities. In figure 6D, the overwhelming number of TipDC transcripts corresponding to glycolytic proteins were increased. Likewise, most of the proteins in the glycolytic protein network from in vitro IAV infected DC were increased (S1D Fig). In terms of glycolysis, TipDC significantly increased hexokinase, phosphofructokinase, triosephophate isomerase, phosphoglycerate kinase, and enzymes that convert pyruvate to acetyl CoA and lactate (Fig 6D). We then overlaid gene symbols identified from the TipDC transcripts and in vitro differentiated DC proteins on the enzymes of Glycolysis, pyruvate metabolism, and TCA cycle (Fig 7A). The glycolytic enzymes tended to have positive log2 ratios, indicating an increase with IAV infection, while the TCA cycle enzymes were skewed negatively (Fig 7A). Then we combined the transcripts from each metabolic pathway compared them. We found the IAV to control log2 ratios of transcripts of glycolytic enzymes were significantly higher than those of the TCA cycle for in vitro differentiate DC (Fig 7B). Likewise, when we performed the same analysis of the TipDC isolated from mouse lungs following IAV, we found a very significant difference in IAV to control log2 ratios of glycolysis verse TCA cycle enzyme transcripts (Fig 7B). In both these cases, the values above zero log2 ratio reflect enzymes whose gene transcripts or protein peptides increased with IAV. Thus, irrespective of strain used in vitro, in vitro or in vivo DC differentiation, or infection in vitro or in vivo we find dendritic cells globally reprogram metabolism in response to influenza infection.

**Figure 7.**
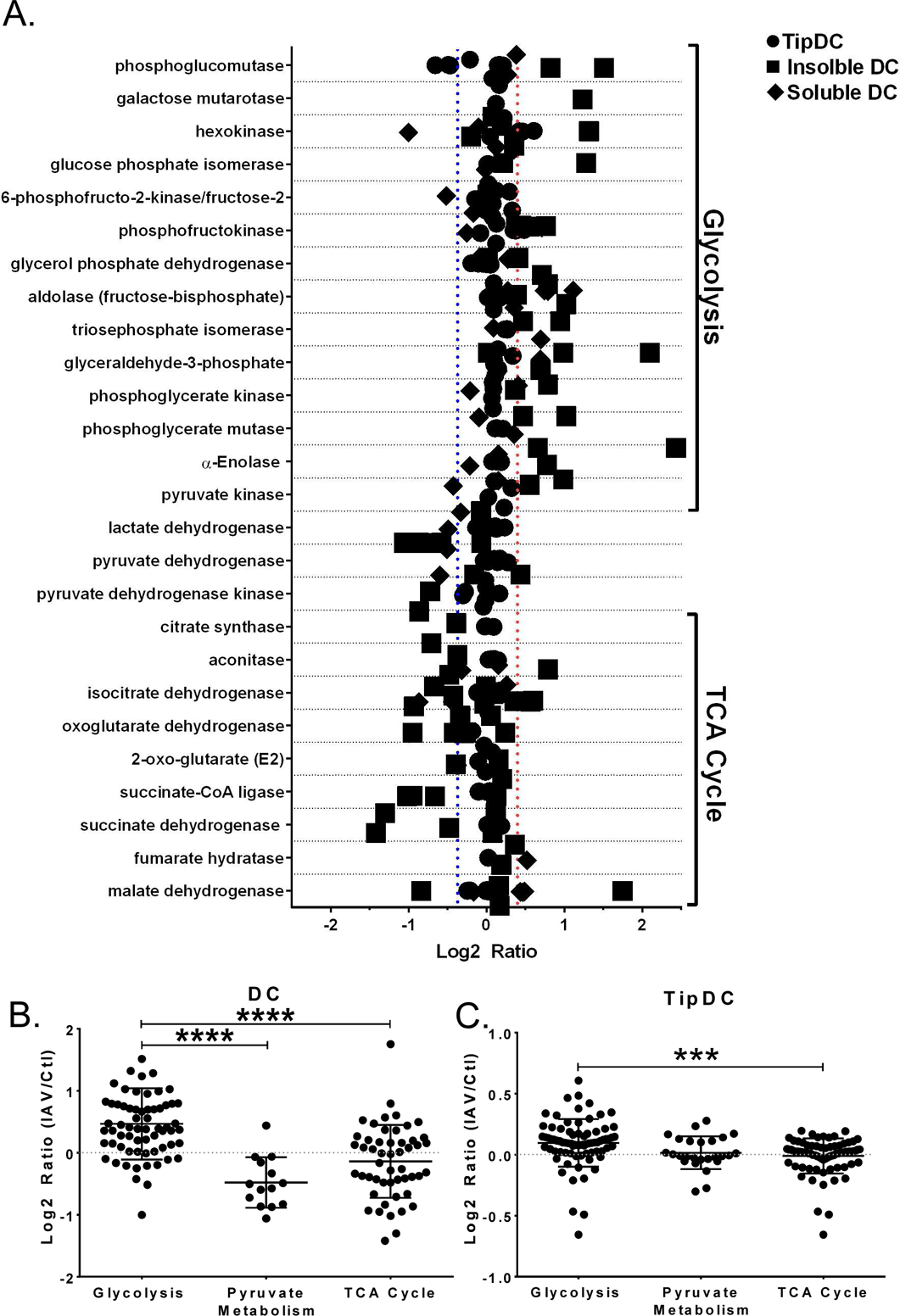
Dendritic cells globally reprogram metabolism in response to IAV infection. Proteins were extracted from in vitro differentiated uninfected and infected DC, separated into insoluble and soluble fractions and subjected to mass spectrometry. RNA was extracted from lung TipDC on day 0 or 9 of IAV infection and subjected to RNA-Seq. The log2 ratio (IAV/Ctl) of transcripts and peptides were determined. Identifiers were converted to gene symbols and official protein names, each dataset were manually searched for all NCBI listed genes and synonyms corresponding to the major regulatory enzymes of glycolysis, pyruvate metabolism, and TCA cycle. (A) All isoforms of each enzyme were grouped (y-axis) and log2 ratios plotted. (B) Transcripts were grouped into glycolysis, pyruvate metabolism, and TCA cycle. (C) Peptides of enzymes were grouped into glycolysis, pyruvate metabolism, and TCA cycle. Graphical abstract. Schematic of global metabolic changes in IAV infected DC. Arrows indicate flow of metabolic products. Bar graph represents the percent difference between control uninfected and infected DC for experiments detailed in Figs 2 & 3 +/- SD.

## Discussion

Our data indicate dendritic cells alter their metabolism in response to IAV infection similar, yet distinct from, our previous findings in human primary respiratory epithelial cells[1]. Unlike our previous observation in epithelial cells, the only significant metabolic difference (p= 0.0102) in DC infected with replicating and non-replicating virus was lowered proton leak (Fig 3C and S3C Fig). This makes sense given IAV non-productively infects DC whereas epithelial cells are the primary source of IAV propagation in the lungs and viral propagation requires high metabolic activity [79, 93, 94]. In our previous epithelial study, we found glycolysis was significantly lower in the absence of viral replication and epithelial cells infected with replicating IAV lack metabolic flexibility and die rather than adapt to changes in available nutrients [1]. In contrast, here we found IAV infection increased DC glycolysis as well as capacity and flexibility for fueling respiration without loss of viability in substrate limiting conditions (S4C Fig). These data reinforce the idea that metabolic reprogramming of parasitized cells reflects aberrant changes driven by viral replication while infected dendritic cells are harboring the virus while maintain metabolic control. However, we found processes such as non-glycolytic extracellular acidification, non-mitochondrial oxygen consumption, and glucose uptake were significantly lower when IAV was BPL inactivated. These differences in DC transduction of signals that govern metabolic substrate import and product export may be worthy of closer examination.

Viral infections, like those caused by IAV, have devastating consequences for cancer patients, who often receive metabolic modulators in their treatment regimens that have the potential to alter immune function. Our current findings, in the context of our previous work, indicate the effects of these treatments may be cell and pathogen specific. Indeed, in the past 10 years there has been considerable focus on shared pathways in T lymphocytes and cancer metabolism and more recently it has become clear that APC metabolism can also be altered by activation with TLR agonists PMA and LPS [14, 16-19, 41, 95, 96]. However, little was known about the metabolic demands of innate immune cells in response to IAV infection or IAV associated TLR activation. Based on the work of pioneers in this field, we anticipated IAV and LPS would have stark differences in bioenergetics [18, 19, 59, 61, 97]. Indeed, when we compared IAV infection to LPS activated DC we found basal respiration, maximal respiration, coupling efficiency, and ATP production were significantly increased (Fig 3C and S3C Fig). Perhaps the most striking and significant difference was in glycolytic ATP production, where IAV infected DC produced an average of 393.5 pmol/min/RFU 95%CI [259.9, 527.1] more ATP than LPS treated DC (Fig 3D and Sup3D). Using both healthy controls and LPS as reference points, allowed us to compare a spectrum of mitochondrial fitness and determine that IAV infected DC retain respiratory function based on their FCCP response in the mitochondrial stress test (Fig 3). Further, while the proportion of ATP production from OXPHOS was significantly lower than that produced by glycolysis, the mitochondrial ATP production upon IAV infection was only reduced by 20% in DC indicating that, unlike with LPS treatment, respiration was not severely impaired. However, we were surprised to find DC stimulated by TLR agonists with considerable overlap in signaling pathways (i.e. endosomal TLR3 and 7/8 viral RNA mimetics PolyIC or R848, respectively) had a very different response to FCCP. This resulted in significant differences in maximal respiration, spare respiratory capacity, coupling efficiency, and mitochondrial ATP production (S3C Fig). The later was confirmed with an FCCP independent ATP rate assay that showed R848 produced 57.47 pmol/min/RFU 95%CI [-75.56, −39.38] less mitochondrial ATP than PolyIC (p < 0.0001) even though both agonists increased DC glycolysis similarly (Fig 3A&D and S3D Fig). Thus, we have highlighted an additional layer of complexity in the mechanisms underlying activated DC metabolic rewiring and found that TLR agonists and IAV infections induce unique bioenergetics. Future studies in this area will better dissect these signaling pathways and tease apart how they orchestrate metabolic reprogramming.

Using quantitative proteomics and transcriptomics we found global rewiring of metabolism early in the DC response to IAV infection in vitro and in TipDC 9 days following IAV infection in vivo. We found a pronounced shift in metabolic protein abundance levels and localization in DC resulting in significant enrichment of metabolic pathways, and this was validated by measuring metabolism of specific substrates, extracellular and intracellular metabolite concentrations, substrate product and flux rates for glycolysis and respiration, and bioenergetic functional assays of ATP production or metabolic capacity and flexibility. Combined these observations delineate the central features of the metabolic response of DC to IAV infection, showing a significant increase in glycolysis and glycolytic ATP production, that is decoupled from canonical pyruvate oxidation in the TCA cycle in favor of utilizing glutamine. This phenomenon is reminiscent of

Warburg metabolism, a phenotype that reflects an increase in demand for anabolic metabolites and the energy to assemble them into macromolecules necessary for cell proliferation. Further, this metabolic reprogramming has been shown to be pivotal for proliferation and effector functions of other immune cells [14, 15, 39-42, 84, 85, 96, 98]. Given IAV infected DC have increased capacity to utilize diverse substrates and the flexibility to toggle between them, we are currently unable to fully explain why inhibiting oxidation of specific substrates in the mitochondria would significantly reduce migration or effect DC priming of T cells (Fig 2&5). However, these mitochondrial inhibitors did not ablate the DC response to infection, and this may indicate the metabolic compensation by DC to use suboptimal mitochondrial fuels comes at a cost that diminishes their effector function. Our findings extend our understanding of immune metabolism into influenza infection of a critical cell type that bridges innate to adaptive immunity and establishes an appropriate in vitro model to study of this metabolic phenomenon.

It has long been appreciated that DC migration from the lungs to draining lymph nodes is critical to mounting an effective immune response to IAV, and motility is an energy-demanding process that is directly coupled to ATP availability [79, 90, 99, 100]. We observed a dramatic increase in DC mobility following IAV infection and found IAV-infected DC derive approximately 17 times more ATP from glycolysis than mitochondrial OXPHOS corresponding to 10 fmol/min verses 0.6 fmole/min per infected DC (Fig 5A and 3D). One intriguing possibility is that this pool of glycolytic ATP is critical for infected DC trafficking. In support of this, we found inhibiting c-Myc abolished IAV induced glycolysis and translocation of DC (Fig 4B and 5A). IAV-infected DC appear to acquire a high level of plasticity in energy substrate metabolism, including greater carbon and energy source flexibility. Indeed, decoupling of glycolysis and TCA cycle in cancers was once presumed to be a mitochondrial defect but is now postulated to produce a more robust and flexible metabolic program [43, 101]. c-Myc is considered a master regulator of cancer metabolism, and metabolic reprogramming by c-Myc is thought to allow cancer and activated immune cells to sustain supplies of anabolic building blocks while generating energy for their assembly[14, 63, 64, 68, 72]. Our data supports applying this idea to IAV-infected DC, given we found c-MYC expression increased in response to IAV infection and c-Myc inhibition significantly reduced glycolysis, respiration, and glutamine import, while preventing IAV-induced DC migration and induction of CD25+ and CD44+ CD8 T cells (Figs 4, 5A-F, and S5E Fig). However, an important and related recent discovery is that, as DC mature they down regulate expression of c-MYC and n-MYC while maintaining constant expression of l-MYC even after activation [68, 69]. Paradoxically, mice with l-MYC-deficient dendritic cells have reduced T cell response during either bacterial or viral infection [68, 69]. In this respect, while it is possible that our reagents were nonspecific we suggest that is unlikely for several reasons: the primers we used are specific for c-MYC [102], these primers only amplified one species, and the c-Myc antibodies we used detected a single band at the appropriate size. Moreover, we observed a transient increase in response to activation stimuli (i.e. IAV infection) at the RNA and protein levels, whereas others report l-MYC remains constant in activated DC [69]. Importantly, we found c-Myc inhibitor treatment blocked or diminished critical DC immune functions. The exact cause of this effect remains unclear, but it correlates with a loss of DC glycolytic response to IAV. With respect to the interplay of the Myc paralogs and determining their specific roles in relationship to DC stimuli, phenotype, and pathogen response there remains an open questions [69, 103] that now include the specific role of c-Myc in IAV infection.

In addition to its other roles in cancer metabolism, c-Myc directly regulates glutaminase, transcriptionally activates glutamine transporters, promotes glutamine metabolism including glutamine catabolism to glutamic acid for anaplerosis and NADPH, and in cancer c-Myc transcriptionally coordinates increased glutaminolysis, dependence on exogenous glutamine and glutamine addiction that is c-Myc dependent and can’t be rescued by anti-apoptotic proteins or TCA intermediates[65, 66, 104-109]. In cancer, glutamine utilization confers several advantages over classical glycolysis coupled to TCA cycle for energy production. This is primarily because-ketoglutarate, and steps that follow it, normal levels of reducing equivalents are produced independently of Acetyl CoA, glycolysis, and FAO while supplying ample building blocks for de novo protein synthesis, purine nucleoside biosynthesis, and de novo fatty acid and cholesterol biosynthesis. Thus, glutamine metabolism has the added benefit of increasing proton buffering capacity and cellular redox homeostasis including glutathione synthesis from glutaminase enzymes [71, 110] and when glutaminolysis is inhibited ROS increases [111]. Meanwhile, the first step of the TCA cycle is sensitive to ROS [64, 112-114] as is the respiratory chain and membrane permeability, and this mitochondrial damage potentiates ROS [115, 116]. Hence, it is conceivable that glutamine oxidation may have important advantages for an activated innate immune cell that produces ROS in conjunction with other effector functions that require high anabolism and energy production. This is consistent with our observations that infected DC redirect a large portion of pyruvate away from the TCA cycle while simultaneously increasing glutaminolysis, capacity for oxidizing glutamine, glutamine import and significantly depleting intracellular glutamine concentrat-ketoglutarate concentrations if the mitochondrial pyruvate carrier in inhibited and increase c-Myc (Fig 2C&D, 4A, and S2E Fig). Recently sustained import and utilization of glucose and glutamine were shown to stabilize c-MYC and were required for T cell renewal and clonal expansion [117]. Determining the mechanistic underpinnings of glutamine utilization in dendritic cells will advance our understanding of their basic metabolism and will lead to a clearer picture of immune function in the IAV-infected lung microenvironment.

In conclusion, our findings support the view that dendritic cells specifically rewire their metabolism based on activation stimuli or IAV infection and the latter significantly increased glycolysis, glycolytic ATP production, glutaminolysis, and metabolic plasticity concomitant to reducing pyruvate utilization in the TCA cycle. This metabolic reprogramming is preceded by a transient increase in c-Myc expression and ablating its activity restored normal glycolysis and motility and reduced T cell priming. We clearly demonstrated IAV infected DC acquires the majority of ATP from glycolysis, nevertheless restricting mitochondrial fuels significantly impairs DC function. Collectively our data indicate increased metabolic rewiring and plasticity while also highlighting the impact of specific metabolic substrates on DC migration and T cell priming. This is significant, given reducing DC migration after viral infection profoundly reduces the amplitude of the T cell response and the strength of antigenic stimulation regulates T cell progression [90, 118]. It remains to be determined if these restraints on DC metabolism result in a defect in antigen capture, processing or MHC loading/presentation, or if long term quantity or quality of expanded virus-specific T cells is impacted. Importantly, it appears dendritic cells orchestrate coordinated bioenergetic and metabolic changes in response to IAV infection irrespective of strain and are similar regardless of DC origin or infection model (i.e. in vitro differentiated or lung Tip DC and in vitro or in vivo, respectively). Although incomplete, these data suggest the net result of blocking metabolic reprogramming therapeutically in patients or limiting metabolites in situ could have detrimental consequences in terms of mounting an appropriate immune response. It is also tempting to postulate that local metabolite depletion in the lung may act to limit lung injury by placing a governor on the immune response. However, the metabolite milieu of the IAV-infected lung and lung-lining fluid has not been well established, and this must be addressed to determine if variability in metabolite concentrations impact cellular responses or secondary infections. It also remains to be determined if metabolic reprogramming occurs in humans with community-acquired respiratory viral infections and, if so, if it can be exploited therapeutically to enhance or reduce the immune response to IAV.

## Materials and Methods

### Ethics Statement

All experiments using mice were done in compliance with the Guidelines of Care and Use of Laboratory Animals and approved by University of Tennessee Health Science Center Animal Care and Use Committee. Experiments were conducted under IACUC protocol # 16-138.0.

### Tissue Culture and Mouse work

#### Bone marrow derived dendritic cells (DC)

Female C57BL/6J (B6) mice purchased from The Jackson Laboratory (Bar Harbor, ME) and were maintained in specific-pathogen-free facilities at the University of Tennessee Health Science Center (UTHSC), Memphis, TN. Primary bone marrow derived dendritic cells were isolated and differentiated as previously described (104). Briefly, after removal of the epiphysis bone marrow was flushed out, red blood cells lysed, cell viability determined with AO/PI and 2 million viable precursors per 100 cm^2^ polystyrene plate were seeded in 10 ml RPMI supplemented with Penicillin (100 U/ml), Streptomycin (100 µg/ml), fetal bovine serum (10% v/v) (Gibco Grand Island, NY) and 20 ng GM-CSF (R&D systems, Minneapolis, MN) per ml medium. 20 ng GM-CSF/ml medium was supplemented on days 3, 6, and 8 then DC were harvested on day 10.

#### T cell isolation

Mice were chemically restrained with 2,2,2-tribromoethanol (Sigma Aldrich) with a dose adjusted to the individual body weight and infected intra-nasally with 10^4^ LD_50_ of PR8 dose over a 21-day period. Mice were sacrificed and spleens harvested from virus challenged (primed) or naïve 8-12 week-old C57BL/6 or C57BL/6-Tg(TcraTcrb)1100Mjb/J (OT1) male and female mice. T cells from challenged mice were only used for flow cytometry experiments. Spleens were manually disrupted by grinding organ tissue between the frosted ends of two sterile glass microscope slides in sterile PBS containing 2% fetal bovine serum. T cells were isolated by magnetic separation using EasyStep^TM^ Mouse CD8+ T Cell Isolation Kit with anti-CD8 magnetic beads (StemCell^TM^ Technologies) (purity>99%) for CD8+T Cells or EasySep™ Mouse T Cell Isolation Kit (StemCell^TM^ Technologies) for T cells isolation by negative selection.

#### Intranasal infection and lymphocyte isolation

Female and male 6–8 week-old C57BL/6 mice were purchased from Jackson Laboratories (Bar Harbor, ME). All mice were acclimated for two weeks prior to inoculation with IAV. All infections were conducted in an ABSL2+ facility where animals were assessed daily. Those with severe morbidity (greater than 30% weight loss plus severe clinical impairment) were humanely euthanized according to our approved protocol. Prior to EID_50_ 2000 the mice were chemically restrained by intraperitoneal injection of 2,2,2-tribromoethanol (Avertin). Throughout the course of the infection mice were monitored by daily weighing and assessment of the clinical distress symptoms (e.g. ruffled fur, hunched back, lethargy etc.). At day 0 or 9 mice were euthanized and the lungs were collected after perfusion in situ via the ventricles with 40–50 ml of PBS. The lungs were rinsed with PBS, minced with fine scissors and gently pushed through a 40-syringe plunger. The suspension was then washed with Click’s medium and incubated in digestion buffer (Click’s plus 50 U/ml collagenase IV and 0.001% DNase I (Sigma-Aldrich)) for 30-90 minutes at 37°C on a rocking platform. Lungs tissue was then passed through a 1µm sieve and cells collected, washed (40 ml of Click’s medium), and resuspended in sterile 2%-PBS. Lymphocytes were isolated with percoll density gradient.

#### Virus propagation and titer

A/PuertoRico/8/34, referred to in the text as IAV, is a mouse adapted highly pathogenic influenza virus (PR8) [119]. This virus was cultured in pathogen free antibiotic treated eggs (generously provided by Dr Paul Thomas) that was inoculated on day 9 in bulk from a single viral stock each. Allantoic fluid was harvested 48 hours following inoculation and aliquoted into 50 ml conical tubes and stored at −80°C. Individual bulk viral stock was then thawed, aliquoted, stored at −80°C and titer was determined. Tissue culture infectious dose at 50% (TCID_50_) were then determined by serial dilution on near confluent Madin-Darby canine kidney (MDCK) cells (purchased from American Type Culture Collection (ATCC CCL-34)) in the presence of 1 mg/ml trypsin (Sigma Aldrich), verified with 50% egg infectious doses (EID_50_) and calculated according to the Reed and Muench method [120]. A/PuertoRico/8/34 virus was added at a multiplicity of infection (MOI) of 5 for 2 hours to DC. Virus laden medium or blank infection medium was then removed, and the infection was allowed to proceed for indicated times.

#### Proteomics: sample preparation, mass spectrometry, and data processing

We have previously described these proteomics methods elsewhere in detail. Details regarding sample acquisition, MS parameters, and data processing can be found in Smallwood et al 2011 (PMCID: PMC3241613). Details specific to quantitative proteomics using isobaric tags can be found in Smallwood et al 2017 [1]. Details specific to quantitative proteomics using stable isotopically labeled proteomics see Lopez-Ferrer et al 2009 [23]. Brief summaries of pertinent details are also provided below.

#### Sample Preparation

DC were infected for 17 hours with PR8 (MOI 5 or 1) followed by lysis, homogenization, and fractionation by centrifugation. iTRAQ samples were tryptically digested by using a FASP Protein Digestion Kit (Protein Discovery, San Diego, CA) according to the manufacturer’s instructions with slight modifications (Supplemental Data), samples were desalted by C18 solid-phase extraction (SPE) before isobaric labeling (SUPELCO). Digested samples were then processed according to the manufacturer’s directions for iTRAQ 4-plex labeling (ABSciex). A separate set of DC samples was processed by performing trypsin-catalyzed ^18^O labeling. Before fractionating via high-pH reverse-phase fractionation with concatenated pooling, samples were desalted by C^18^ SPE (SUPELCO). All samples were processed in a custom liquid chromatography system using reversed-phase C^18^ columns. The iTRAQ samples were analyzed by using a Velos Orbitrap mass spectrometer (Thermo Scientific), and ^18^O-labeled samples were analyzed by using an LTQ-Orbitrap mass spectrometer (Thermo Scientific). Both systems were equipped with custom ion funnel–based atmospheric pressure ionization sources and electrospray ionization interfaces.

#### Data processing

Raw files were compared with a concatenated NCBI Mus musculus database and contaminant database by using SEQUEST v.27 (rev. 12). The resulting sequence identifications were rescored by using MS-GF and filtered to a 1% false-discovery rate by using the target-decoy approach and MS-GF–derived spectral probabilities. Reporter-ion intensities were quantified by using the MASIC tool (2). Missing reporter ion channel results were excluded from analysis. Redundant peptide identification reporter ions were summed across fractions, and median central tendency normalization was used to account for channel bias and then log_2_- transformed. Soluble and insoluble fractions were analyzed separately by using DanteR software. The analysis of variance model included treatment and peptide effects. The remaining treatment effect and p-value were calculated by using Student’s t-test. Benjamin-Hochberg multiple-testing error correction was applied to the results of this hypothesis testing, and proteins having a p-value 0.05 were considered statistically significant.

#### Bioinfomatics

Peptide identifiers were converted to ENSEMBL and ENTREZID gene identifiers and protein names with DAVID. The log2 ratio of statistically significant peptides was converted to fold change and the soluble and insoluble proteins were separated by fold change (i.e. increasing and decreasing 2-fold). iTRAQ and SIL proteomes were combined, and redundancies removed by selecting the most significant peptide. The 2-fold protein lists were then submitted to DAVID for enrichment analysis as previously described [24]. KEGG Orthology (KO) pathways were then cross referenced with KO Class and subcategory. Network summary of localization changes in confidently identified proteins with significant enrichment in metabolic pathways was performed as previously described [25]. Briefly, subclasses of KO metabolic proteins were organized based on location in the soluble or insoluble fraction and connected to protein nodes with red or blue edges based on increasing or decreasing abundance, respectively. Protein nodes were then colored by KEGG subclass: carbohydrate metabolism (red), amino acids metabolism (orange), nucleotide metabolism (purple), energy metabolism (green), lipid metabolism (yellow), and glycan biosynthesis and metabolism (blue). Cytoscape network was built essentially as previously described [121]. Briefly, we submitted both the soluble and insoluble SIL proteomes to DAVID to select proteins in the KEGG glycolysis pathway. We used PPI spider to determine the glycolytic protein-protein interaction network [122]. We then mapped the interaction network to the glycolytic network and used Cytoscape to overlay abundance data from the proteomes.

### Transcriptomics: sample preparation, RNA Seq, and data processing

#### Sample preparation

RNA was extracted from sorted Tip DC using 1 mL TRIzol reagent (Invitrogen). RNA was extracted from Trizol suspensions using the Zymo Research Direct-zol RNA miniprep kit, treated with 10 U DNAse for 20 min at RT and then concentrated using the Qiagen RNeasy MinElute clean up kit. Purified RNA quality checked using the Nano Total RNA kit on an Agilent 2100 Bioanalyzer.

#### RNA seq

We used Illumina’s TruSeq RNA v2 sample preparation protocol according to manufacturer’s instructions for generation of RNA-seq libraries. Briefly, this protocol involved the following steps: A. cDNA Synthesis: mRNA was purified via oligo(dT) beads followed by fragmentation using divalent cations and heat. The 1st strand cDNA synthesis was performed using random primers, followed by 2nd strand cDNA synthesis. B. cDNA library preparation. cDNA fragments were DNA blunt end repaired followed by addition of 3’ adenylation of DNA fragment. Sequencing adapters were then ligated to the ends utilizing T-A pairing of adapter and DNA 2 fragments. This was then followed by PCR amplification of the library. The cDNA libraries were run on an Illumina HiSeq2000.

#### Data Processing

Data quality was checked with fastqc software. Paired end RNA-Seq data was aligned to Mouse mm10 genome using TopHat (with Bowtie2). Only concordant pairs with mapping quality > 10 were kept. The number of reads assigned to each gene was found using Bioconductor R package. Count data was analysed using edgeR Bioconductor package (GLM formulation). Before doing this genes, which did not have >= 3 counts in every sample for at least one group, were filtered out. Genes were declared DE if they had FDR < 0.05 and log2 FC > 1. Multidimensional Scaling (MDS) as implemented in EdgeR function plotMDS (in two dimensions) was used for data visualization. Briefly, MDS takes as input the set of pairwise distances between any two samples and plots points in 2-dimensional space (plane) attempting to preserve as much as possible the original distances between any two samples.

#### Bioinfomatics

Transcript identifiers were converted to official gene symbols with DAVID and linked to expression levels in Excel. The XLSTAT OMICS data analytics package was used for generating the heat map and expression analysis. Differential expression was determined from three day 0 controls and four day 9 IAV infected samples. The standard deviation threshold for this analysis was 50%, eliminating 6,643 transcripts out of 13,285. Significant differences in expression were selected based on p-values generated from a parametric comparison of differences using Tukey honest significant difference test for multiple comparisons with Benjamini-Hochberg post-hoc corrections.

### Metabolic assays

#### Metabolite depletion – Live/dead viability assay

DC were cultured as described above. After 24 hours of adherence, RPMI 1640 complete medium was replaced with fresh RPMI 1640 complete medium or RPMI metabolite depleted medium for 3 hours (i.e. RPMI without glucose or without glutamine or without both, respectively). Later the cells were treated as above in complete or deplete media followed by infection for 17 hours. The cells were labelled with calcein-AM (2µM) and EthD-1(4µM) incubated for 20 minutes at room temperature and the viability was measured using Live/Dead Viability/Cytotoxicity assay kit for mammalian cells (Thermo Scientific) with a fluorescent plate reader at 494/517nm and 528/617nm, respectively.

#### Metabolic Flux

Glycolytic flux was determined by measuring the detritiation of [3-3H]glucose (Sigma Aldrich) as previously described [123]. Fatty acid beta-oxidation flux was determined by measuring the detritiation of [9,10-^3^H]-palmitic acid (Sigma Aldrich) as previously described [124]. Glutamine oxidation flux was determined by calculating the rate of ^14^CO_2_–release from [U-^14^C]- glutamine (Sigma Aldrich) as previously described [125]. Pyruvate oxidation flux was determined by calculating the rate of ^14^CO2–release from [2-^14^C]-pyruvate (Sigma Aldrich) as previously described [126]. Glucose oxidation flux through the pentose phosphate pathway (PPP) was determined by calculating the rate of ^14^CO2–release from [1-^14^C]-glucose (Sigma Aldrich) as previously described [14, 127]. The concentr using the Free Fatty Acid Assay (AbCam) by a coupled enzyme assay which results in a fluorometric assay with the fluorescent emission/excitation wavelengths 535/587 nm. The resulting fluorometric product was proportional to the free fatty acids present and quantifiable with the palmitic acid standards.

#### Glucose and Glutamine Quantification

DC were treated as above and after 17 hours cells or supernatants were collected alongside blank medium controls incubated together but without cells. Extracellular glucose was quantified with a glucometer against glucose standards essential as previously described [1]. Glucose levels (mg/dl) in the blank medium was determined alongside DC +/- treatments. Glucose uptake was calculated by subtracting the glucose in the control, IAV, IAV^BPL^, PolyIC, R848 or LPS medium from the cell free blank medium. Internal and external glutamine was quantified from supernatant and cell lysates, respectively, following the Glutamine/Glutamate Glo assay manufacturer’s protocol (Promega). Upfront sample processing such as dilution is required to fit in to the linear range of the glutamine or glutamate standard curve. All the standard and sample dilutions were prepared in PBS and for each assay 25 µl sample and 25 µl standard were transferred to a 96 well luminescence assay plate and 25 µl of Glutaminase Buffer was added. The reactions were incubated for 30 minutes at room temperature. Later, 50µl of Glutamate Detection Reagent was added and incubated for additional 60 minutes. The metabolite detection was done using bioluminescent NADH detection method consisting of glutamine conversion to glutamate by Glutaminase and glutamate detection with the Glutamate Detection Reagent (containing glutamate dehydrogenase, NAD+, Reductase, Reductase Substrate and Luciferase) and luminescence was measured using BMG CLARIOstar microplate reader.

#### Bioenergetic assays on the Xfe96 bioanalyzer

ECAR, OCR, and PPR were measured by using the XFe-96 Extracellular Flux Analyzer (Agilent). We first optimized seeding density, plate coatings, and inhibitor concentrations. The final concentrations were: 10 mM Glucose, 50mM 2-DG, 1µM Oligomycin, 2 FCCP, and 0.5 antimycin/rotenone. We essentially followed the manufacture’s protocols. Briefly, 1.5×10^5^ cells per well in Poly D-Lysine coated Seahorse XFe96 microplates at 37°C in 5% CO_2_ for 24 hours. The cells were either left untreated (control) or infected for 17 hours at (MOI) of 5 using viable virus or IAV^BPL^ or treated with LPS (50ng/ml), PolyIC (1µg/ml), or R848 (1µg/ml). At the end of 17 hours of infection the cells were switched to XF base medium (non-buffered DMEM with no phenol red). XF media was supplemented with L-glutamine (2 mM) for Glyco stress test or with glucose (10 mM), L-glutamine (2 mM), and sodium pyruvate (1 mM) for Mito stress test and ATP assay. Injection cartridges were hydrated overnight, cells equilibrated for 1 hour and ECAR, OCR, and PPR monitored for 2 hours. After each run cells per well were quantified with CyQUANT^TM^ (Thermo Scientific) to derive normalization factors. Rates were calculated using the Seahorse XF real time report generator and imported to GraphPad for statistical analysis.

#### Glyco and Mito stress test with c-MYC inhibitor

DC were seeded as described above and were left untreated (control), pretreated with c-Myc Inhibitor (MilliporeSigma) for 4 hrs at 0.5 uM, 1.0 uM, and 2.0uM respectively, infected for 17 hours at (MOI) of 5 with PR8 viable virus.

#### Mito Fuel Flex Test

DC were treated as described above. Calculations were performed with the Seahorse Mito Fuel Flex Test Report Generator plug in for excel and inhibitor injections essentially followed manufacturer protocols using sequential or combined injections of etomoxir, BPTES, and UK5099. Targets and calculations are briefly summarized:

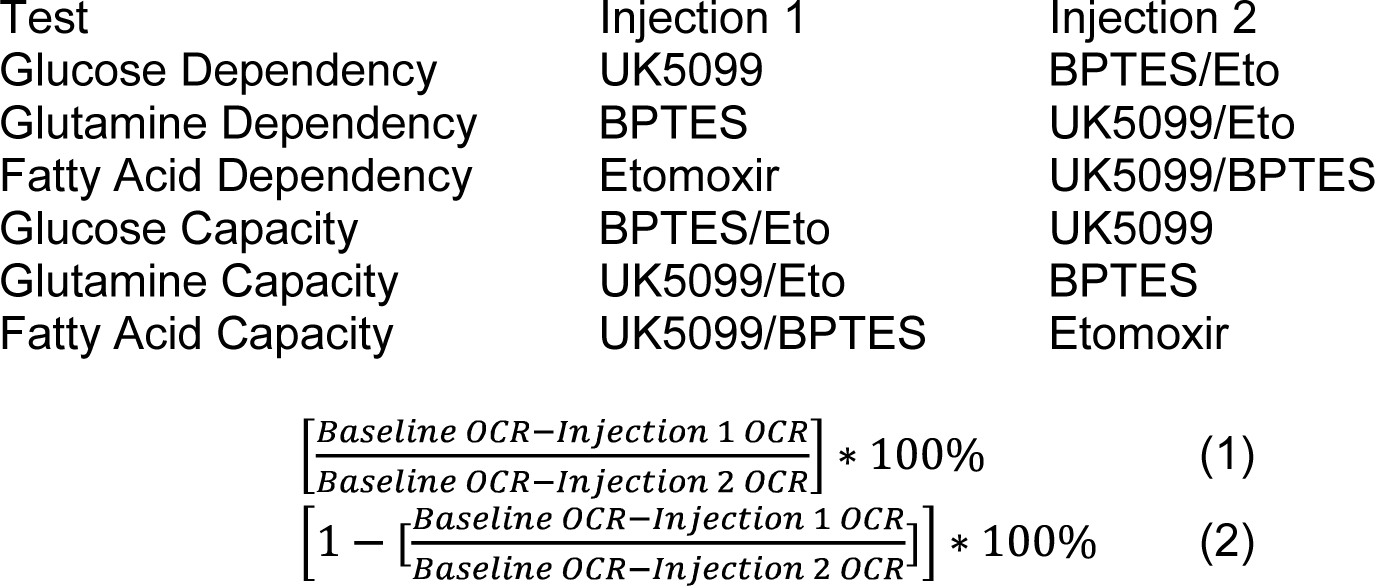

Dependency Equation (1), Capacity Equation (2), and Flexibility (i.e. the difference between the capacity and the dependency for each substrate) were calculated with Seahorse XF Flex Fuel Report generator and imported to GraphPad for statistical analysis. The combined percent oxidation capacity or dependence is the sum of the individual values for glucose, glutamine, and fatty acids.

### Functional Assays

DC were untreated (control) cMyc inhibitor (MI) Bis-2-(5- phenylacetamido-1,3,4-thiadiazol-2-yl)ethyl sulfide (BPTES), UK5099, or etomoxir (Eto) respectively for 4 hours then infected for 17 hours with IAV at MOI of 5. Fresh T cells (purified as described above) were added to DC at 1:5 ratio and incubated for 24 and 48 hours.

#### Cell migration assay

DC were seeded on Poly-D-lysine 100 ug/ml (Sigma Aldrich) precoated Oris™ 96-well migration plate (Platypus Technologies) at the density of 1×10^5^ per well with silicon overnight. After attachment, stoppers were removed, growth media replaced and the DC cells were treated with c-MYC inhibitor (2 uM), BPTES (3 uM), etomoxir (4 uM) or UK5099 (2 uM) and infected with PR8 for up to 17 hours at (MOI) of 5. Following incubation, the cells were rinsed with PBS and labelled with calcein-AM for 20 minutes at 37°C from Live/Dead Viability/Cytotoxicity kit for mammalian cells (Thermo Scientific). After staining, plate was read on BMG CLARIOstar plate reader with FITC excitation/emission filter 485/528nm. The combined effects of migration and stimulation were evaluated by comparing the fluorescent signal detected in the non-infected wells to the signal in the virus infected wells. Data tables from multiple experiments were combined and plotted in GraphPad Prism.

#### FITC-dextran phagocytosis

Cellular uptake of FITC-dextran was determined as reported earlier [128]. DC were left untreated (control) and respectively, infected for 17 hours at (MOI) of 5 with PR8 viable virus and LPS (50 ng/ml) as positive control. After 17 hours of infection cells were labelled with 1 mg/ml FITC-Dextran 40S (Sigma Aldrich) and incubated for 1 hour at 37°C and 4°C, respectively. Uptake was terminated by washing the cells 3x with ice-cold PBS. Cells were resuspended in FACS buffer (PBS containing 2% bovine serum albumin and 0.1% sodium azide). FITC fluorescence of cells was measured using Sony SH800 cell sorter (Sony, Japan). Fluorescence values were reported as mean fluorescence intensity (MFI).

#### Flow cytometry

Fluorescence-activated cell sorting (FACS) analysis was performed using standard methodology. Briefly, after Fc blocking, 1 × 10^6^ cells were stained with appropriate antibodies for 30 min at 4° and then washed with phosphate-buffered saline (PBS)/2% FBS. The cells were stained in FACS buffer (PBS containing 0.5% bovine serum albumin and 0.05% sodium azide) with fluorescently labelled antibodies from BioLegend (CD11c-APC/Cy7, CD11b-BV421, CD40-FITC, CD80-APC, CD86-PE, CD8-PE/Cy7, CD4-PerCp/Cy5.5, CD69-PE, CD25-APC, CD3-BV421, CD44-BD427, CD45.1-FITC, CD3-PerCp/Cy5.5, and CD44-FITC) or Tonbo Biosciences (HLA-DR-PerCp/Cy5.5) to identify DC and T cells. Percentage for each marker was determined by flow cytometry on a Sony SH800 cell sorter and data were analyzed using FlowJo software (Tree Star, San Carlos, CA). Fluorescence minus one (FMO) control for each marker used to identify gating boundaries. DC responses were examined by flow cytometry after staining with anti-CD11c, -MHCII, -CD8, -CD4, -CD11b, -GR-1, -CD103, -CD80, and -CD86 antibodies. TipDC were sorted using CD11b^hi^, Ly6c/GR-1^hi^, MHCII ^hi^ DC surface marker staining on a FACS Aria cell sorter (BDbioscience).

#### T cell activation assay

CD8+ T cells were isolated and co-cultured with drug treated and/or infected DC rate (ECAR) was measured to detect T cell activation in real time using anti-CD3/CD28-coated DynaBeads (Life Technologies) on Xfe96 (Agilent) as previously described [85–88]. ECAR was measured in unbuffered RPMI without phenol red supplemented with glucose (10mM), L-glutamine (2 mM), and sodium pyruvate (1 mM) final concentrations per well. Three rate measurements were done to establish basal rate followed by 15 cycles of real-time measurements of ECAR. Data files were converted to excel files using Wave 2.4.1. Data tables were combined from multiples experiments and plotted and analyzed in GraphPad Prism.

### RNA and protein assays

#### Quantitative Real-time PCR

RNA was isolated from cultured DC and or treated with cMyc for 4, 8 and 17 hours by using Direct-zol RNA MiniPrep Kit (Genesee Scientific) as per the manufacturer’s instructions. cDNA synthesis was performed using the iScript^TM^ cDNA Synthesis Kit (Bio-Rad) per the manufacturer’s instructions. The cDNA synthesis used 7 f reaction mix reaction mix, 1 iScript reverse transcriptase, with the remainder nuclease-free H2O. This was incubated at 25°C for 5 minutes, 42°C for 30 minutes, and 85°C for 5 minutes. Real time RT-PCR was performed using the SYBR^TM^ Green mastermix (Applied Biosystems). Simple relative quantification of target gene expression normalized to actin was performed using the 2 method [129]. The following primers were used: cMyc, forward primer: 5 - CGGACACACAACGTCTTGGAA- - AGGATGTAGGCGGTGGCTTTT-, the primers were purchased from Integrated DNA Technologies.

#### Immunoblotting

Cells were rinsed and pelleted and the pellets were homogenized on ice in a cold Tris-EDTA+0.1% NP-40 lysis buffer. 15 µg of lysates were reduced in NuPAGE reducing reagent (Invitrogen) and boiled for 5 min prior to loading in 4-12% NuPAGE Bis-Tris Gel for separation by electrophoresis. Proteins were transferred to nitrocellulose membrane and blocked with 3% bovine serum albumin (Thermo Scientific) in 1XTBST for 1 hour. The membrane was incubated with cMyc overnight at 4°C followed by washing and incubation with horseradish peroxidase (HRP) secondary antibody. The protein bands were detected using ECL plus (Amersham) on Amersham Imager 600 (GE Healthcare).

## Acknowledgments

Portions of this research were supported by: Le Bonheur Children’s Hospital and the Children’s Foundation Research Institute https://www.lebonheur.org/research-and-education/research/ (H.S.), National Institute of Health National Center for Research Resources (Grant RR018522) https://www.nih.gov/research-training/research-resources (L.P.), The W.R. Wiley Environmental Molecular Science Laboratory: a national scientific user facility sponsored by the U.S. Department of Energy’s Office of Biological and Environmental Research and located at the Pacific Northwest National Laboratory, which is operated by Battelle Memorial Institute for the U.S. Department of Energy under contract DE-AC05-76RL0 1830 (User proposal numbers 37501 & 34745) https://www.pnnl.gov/environmental-molecular-sciences-laboratory (H.S.), National Institute of Health (1UO1CA232488-01 and 1R01AI114581) https://www.nih.gov/ (R.W.), National Institute of Health National Institute of Allergy and Infectious Diseases (R56AI091938 and HHSN266200700005C)) https://www.niaid.nih.gov/ (P.T.), and American Lebanese Syrian Associated Charities (ALSAC) https://www.stjude.org/about-st-jude/faq/whats-alsac.html (P.T.). The funders had no role in study design, data collection and analysis, decision to publish, or preparation of the manuscript. We thank Richard Cross and Greig Lennon for cell sorting and assistance in flow cytometry.

## Supporting information

### Supplemental Figure Legends

**S1 Fig.**
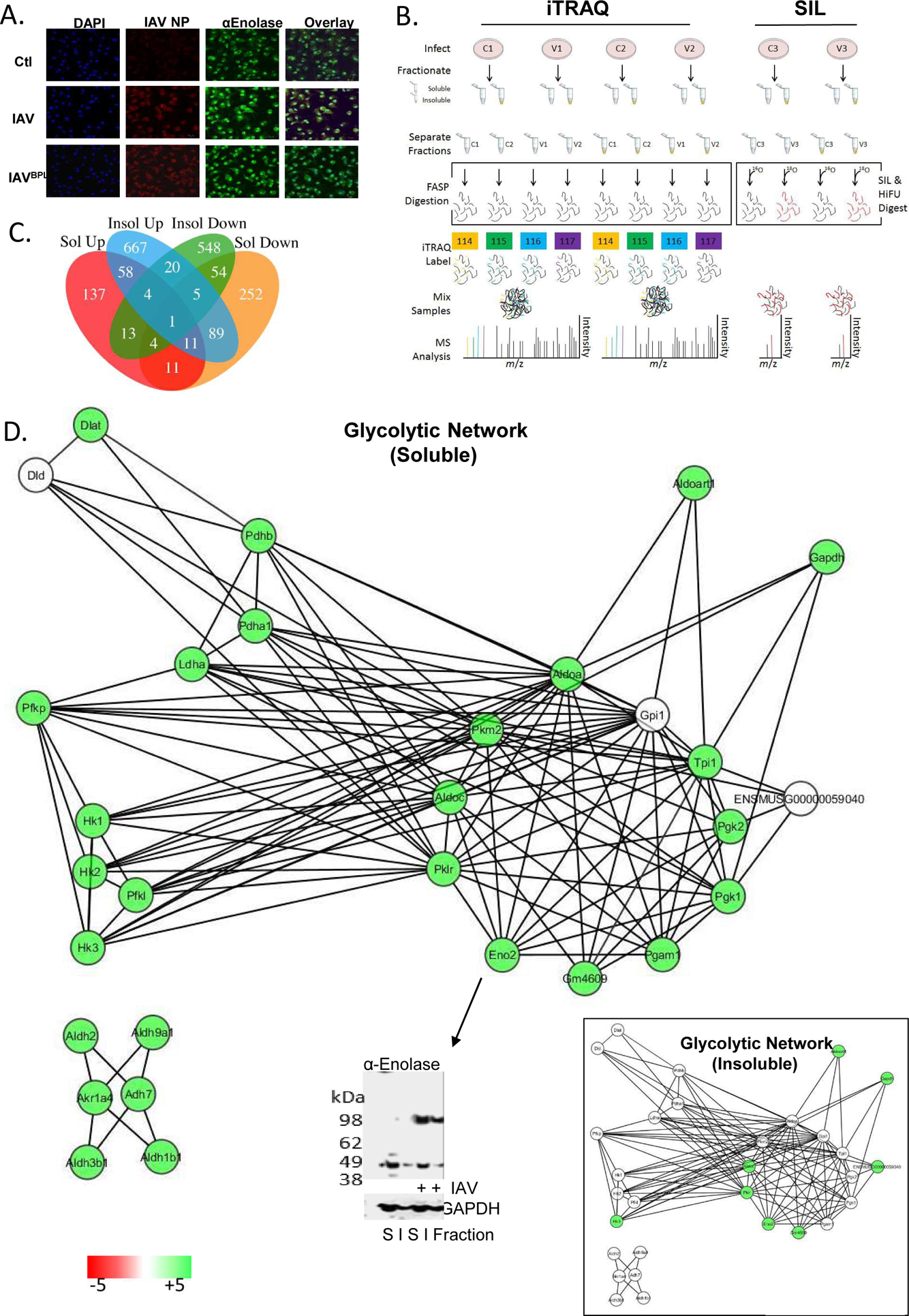
Validation and experimental design of proteomics for IAV infected DC. DC were left untreated (Ctl) or infected for 17 hours at MOI 5 with viable virus (IAV). (A) DC were also infected-propiolactone inactivated virus (IAV^BPL^). DC were fixed and stained for DAPI, influenza nuclear protein or murine-enolase protein and visualized with confocal microscopy. (B) Control uninfected cells or IAV infected BMDC (MOI 5 pfu for 17 hours) were separated into soluble and insoluble fractions. The iTRAQ labeled samples were subjected to FASP digestion while the SIL samples received trypsin-catalyzed ^18^O/^16^O labeling. The samples were desalted with C18 SPE, processed with a custom RPLC system and analyzed with a Velos Orbitrap mass spectrometer (iTRAQ) or LTQ-Orbitrap (SIL). (C) Venn-diagram depicting the distinct total number of proteins identified by iTRAQ and SIL, as well as the overlapping number of proteins. Venn-diagram illustrating the statistically significant number of proteins identified by iTRAQ and SIDL in soluble and insoluble fractions which were up and down regulated as well as the overlapping number of proteins. (D) Both soluble and insoluble SIL DC proteomes were submitted to DAVID and PPI spider to define glycolytic protein-protein interaction networks. The glycolytic network was put into Cytoscape and integrated with quantitative data from the proteomic analysis. Alpha enolase increased in soluble and insoluble (inset) proteomes and was validated with immunoblotting revealing the monomer and dimer increased in both soluble (S) and insoluble (I) networks.

**S2 Fig.**
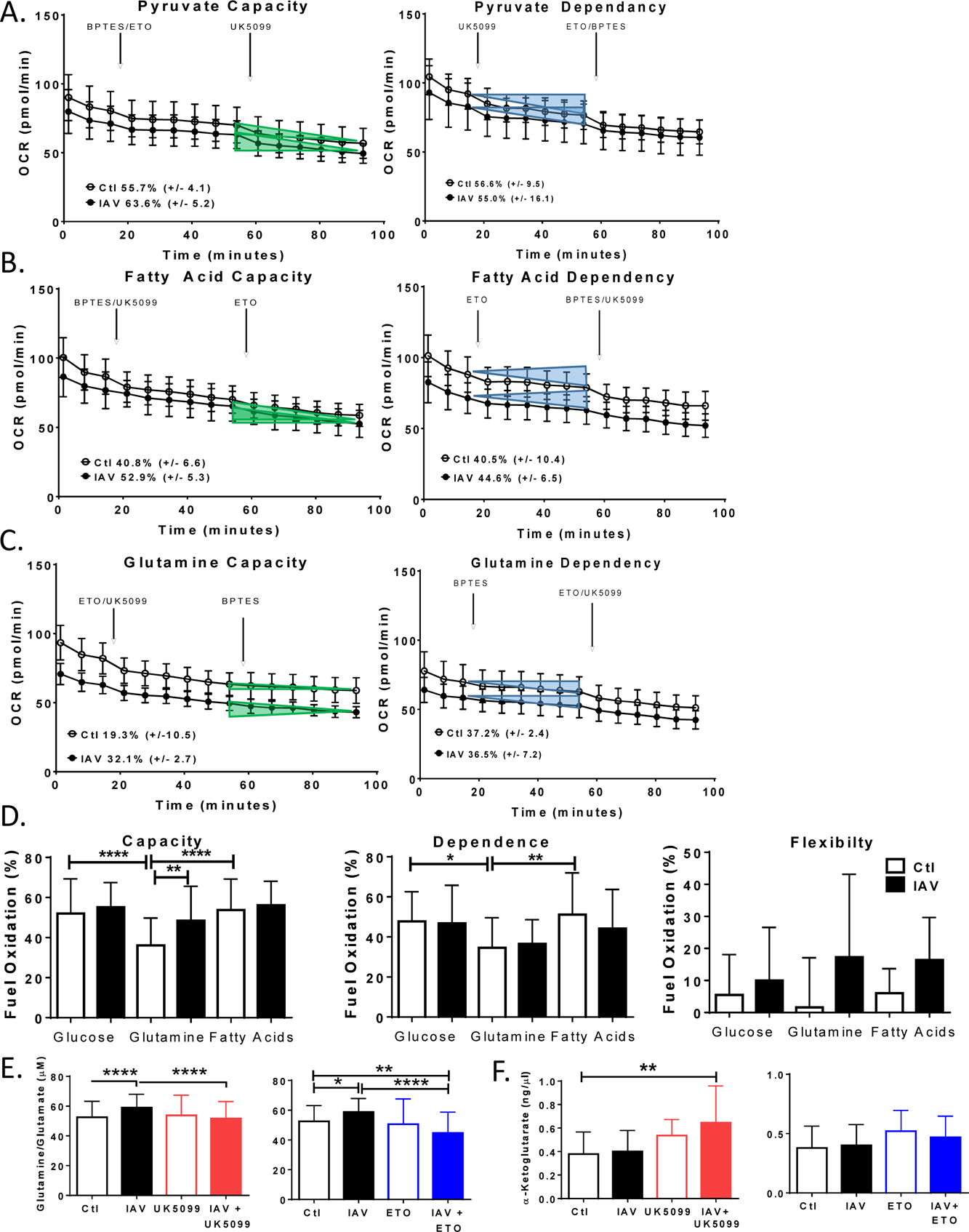
DC metabolic adaptation to fuel respiration following IAV infection. DC were left untreated (Ctl), infected for 17 hours at MOI 5 with viable virus (IAV). (A-D) The rates of pyruvate, glutamine, or long chain fatty acids oxidation for respiration were calculated as the percentage of inhibition of oxygen consumption by UK5099, BPTES, or Etomoxir specific inhibitors of mitochondrial pyruvate carrier, glutaminase, and carnitine palmitoyl-transferase 1A. Capacity for a specific substrate to drive respiratory OCR was tested by determining baseline OCR then inhibiting the 2 off target substrates determining OCR then inhibiting import of the target metabolite. Percent capacity is one minus the baseline OCR less the off-target OCR divided by the baseline OCR less the OCR after all targets inhibited times 100. Dependency on a specific substrate was tested as above reversing the inhibitor sequence and the percent dependence was calculated by deducting the target OCR from the baseline and dividing by the baseline OCR less the OCR after all targets inhibited times 100. Fuel Flexibility was calculated as the difference between Capacity and Dependency. The average capacity of uninfected or infected DC to use either pyruvate, glutamine, or long chain fatty acids was determined. The average dependence of uninfected or infected DC on the oxidation of either pyruvate, glutamine, or long chain fatty acids was determined. The average flexibility of DC to use either pyruvate, glutamine, or long chain fatty acids was determined for uninfected or infected. (E-F) DC were pretreated with UK5099, Etomoxir (ETO), or BPTES +/- IAV for 17 hours, rinsed and lysed for quantification of intracellular glutamine-ketoglutarate activity (F). The bar graphs represent the values of 4-5 independent experiments and presented as experimental mean +/- SD p-value <0.05 (*), p-value < 0.01 (**), and p-value < 0.0001 (****). B-D show one representative experiment of 5 independent experiments with corresponding capacity, dependence, and flexibility values inset.

**S3 Fig.**
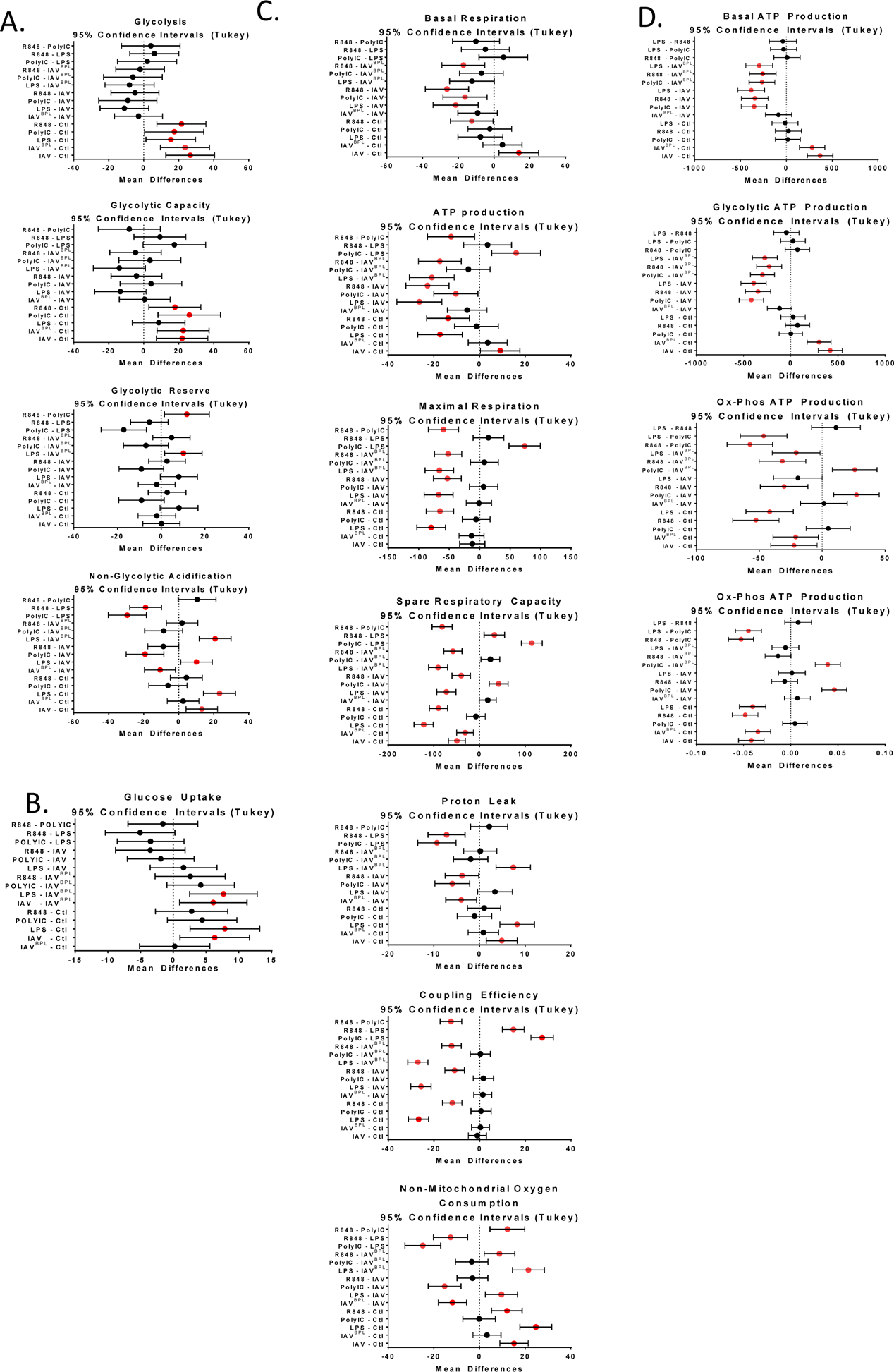
Confidence intervals and mean differences of bioenergetics comparing TLR agonists and IAV. (A-D) DC were infected or treated with TLA agonists lipopolysaccharide (LPS), polyinosinic polycytidylic acid (PolyIC), or Resiquimod (R848) for 17 hours followed by metabolic analysis with Seahorse Xfe96 Flux Analyzer. (A) Glycolytic function was tested while monitoring extracellular acidification rate (ECAR) with sequential injections of glucose, oligomycin (Oligo), and 2-Deoxy-D-glucose (2-DG) indicated by arrows. (B) Glucose uptake was monitored from the medium using a standard blood glucometer with glucose standard calibration curves. (C) Mitochondrial respiration was tested while monitoring oxygen consumption rates (OCR) with sequential injections of oligomycin (Oligo), carbonyl cyanide-p-trifluoromethoxyphenylhydrazone (FCCP), and a mixture of rotenone and antimycin A (Rot/AntA) indicated by arrows. (D) DC maximal mitochondrial ATP changes induced by oligomycin plotted against maximal ATP changes upon glucose depletion determined by respirometry using Xfe96. The graphs represent the difference of the mean values from 3- _replicates_) and 95% confidence intervals. Statistical differences among means was found with ANOVA followed by Tukey, validated with Dunnett’s multiple comparison tests. Dashed line appears at 1 and red circles indicate confidence intervals do not overlap.

**S4 Fig.**
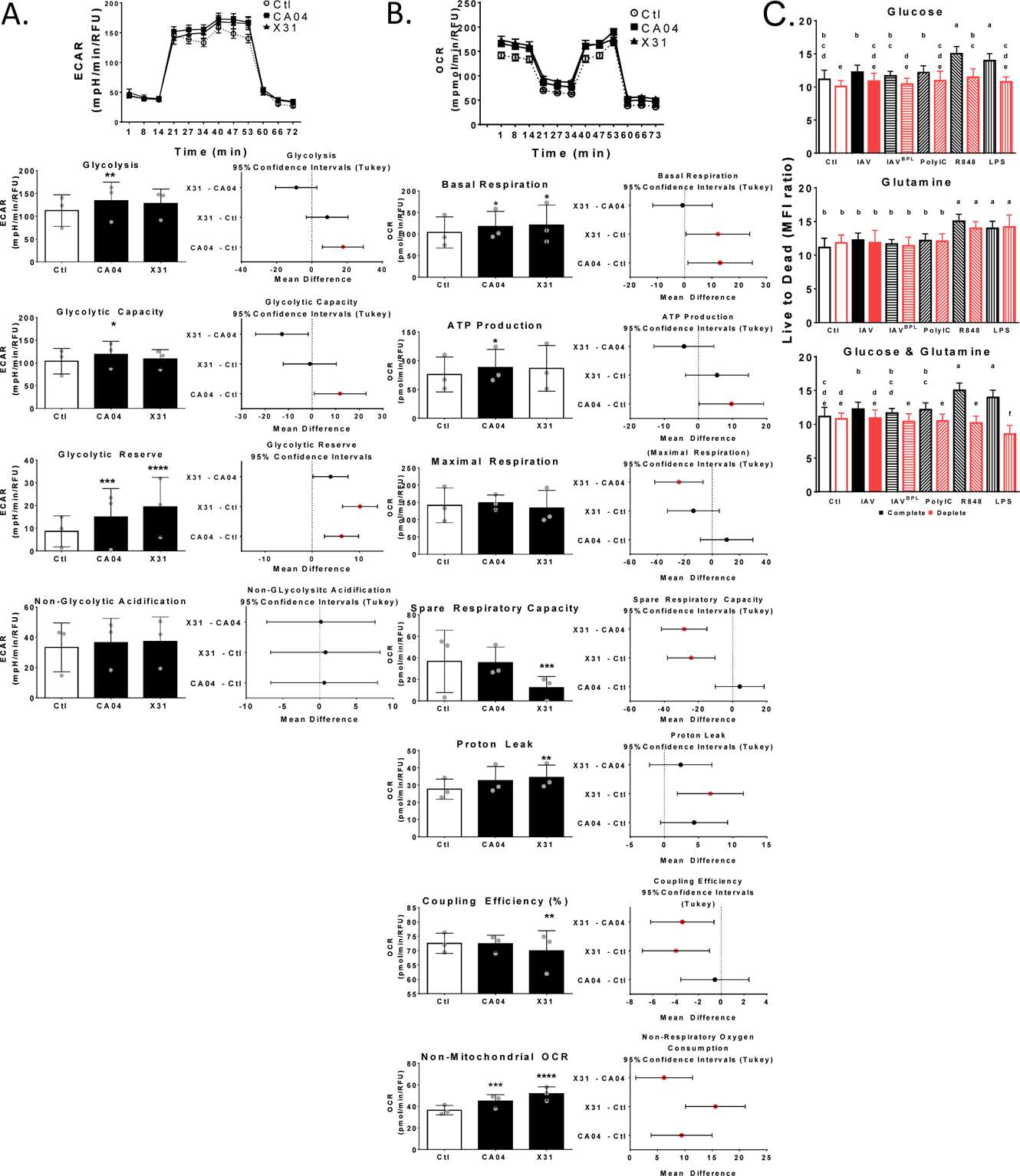
Live to dead ration of DC in response to infection or TLR agonists with metabolite restriction. (A&B) DC were infected with CA04 or X31 influenza virus (MOI = 5) for 17 hours followed by metabolic analysis with Seahorse Xfe96 Flux Analyzer. (A) Glycolytic function was tested while monitoring extracellular acidification rate (ECAR) with sequential injections of glucose, oligomycin (Oligo), and 2-Deoxy-D-glucose (2-DG) indicated by arrows. (B) Mitochondrial respiration was tested while monitoring oxygen consumption rates (OCR) with sequential injections of oligomycin (Oligo), carbonyl cyanide-p-trifluoromethoxyphenylhydrazone (FCCP), and a mixture of rotenone and antimycin A (Rot/AntA) indicated by arrows. The bar graphs represent mean and error bars standard deviation. Statistical differences among means was found with ANOVA followed by Tukey’s Honest Significant Difference (Tukey’s HSD) method. The difference of the means are plotted with Tukey’s HSD 95% confidence intervals. The red circles indicate confidence intervals do not overlap. Tukey adjusted p values are symbolized by asterisks indicating adjusted p-v (C) DC growth medium was replaced with deplete medium 3 hours prior to infection or agonist treatment in blank or viral laden infection medium (MOI = 5) for 2 hours followed by a return to deplete medium for 17 hours. Viable, dead, and total DC were stained with calcein AM, ethidium homodimer and DAPI, respectively, and measured fluorescence intensity using a microplate reader. Then mean fluorescence intensities (MFI) and the ratio of live to dead DC were calculated. The graphs represent the mean values of 3 independent experiments measured +/- SD. Statistical differences among means was found with ANOVA followed by Tukey’s HSD and results summarized using with compact letter display such that groups that are detectably different get different letters.

**S5 Fig.**
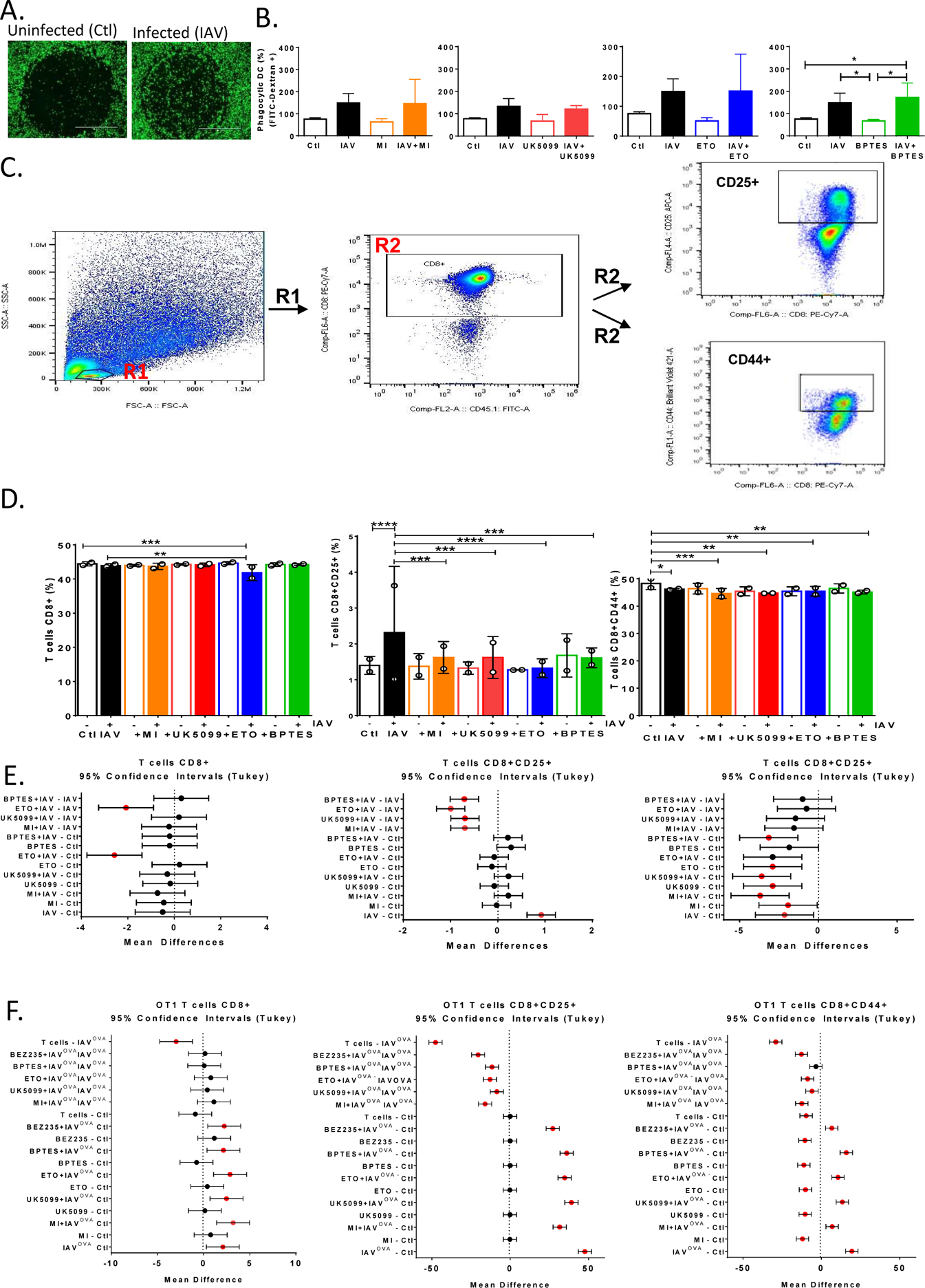
Limiting glycolysis or mitochondrial oxidation of pyruvate, glutamine or long chain fatty acids impairs infected DC function. (A) DC were seeded on precoated Oris™ 96-well migration plate allowed to adhere 18+ hours followed by plug removal and infection (MOI = 5) for 17 hours. Live cells were stained with CalcienAM and images acquired with EVOS, two representative microscopy images of control and IAV migration are presented. (B) DC were left respectively, infected for 17 hours (MOI = 5) with IAV. 1mg/ml FITC-Dextran 40S was added and phagocytosis was terminated after 1 hour at 37°C and 4°C then fluorescence per cell was measured using flow cytometry (C-F) CD8+ T cells were isolated by negative depletion from fresh splenocytes of homologous primed female C57BL/6 mice and co-cultured with DC at 5:1 ratio T cells to DC for 24 hours. Cells were fixed and stained for surface markers then quantified and compensated for using FlowJo software (2 other independent experiments were performed on different instruments and quantified/compensated using different software produced similar trends but were excluded due to compensation methods differences). (C) Representative FACS plots with gating strategy. (D) Bar graphs represent the mean values of 2 independent experiments +/- SD. Statistical differences among means was found with ANOVA followed by Tukey’s HSD with asterisks indicating adjusted p-**** p<0.0001). (E) Tukey’s HSD results summarized with the circles representing the mean difference and the error bars the 95% confidence intervals. (F) Tukey’s HSD results to from Figure 5D are summarized with the circles representing the mean difference and the error bars the 95% confidence intervals corresponding.

**S6 Fig.**
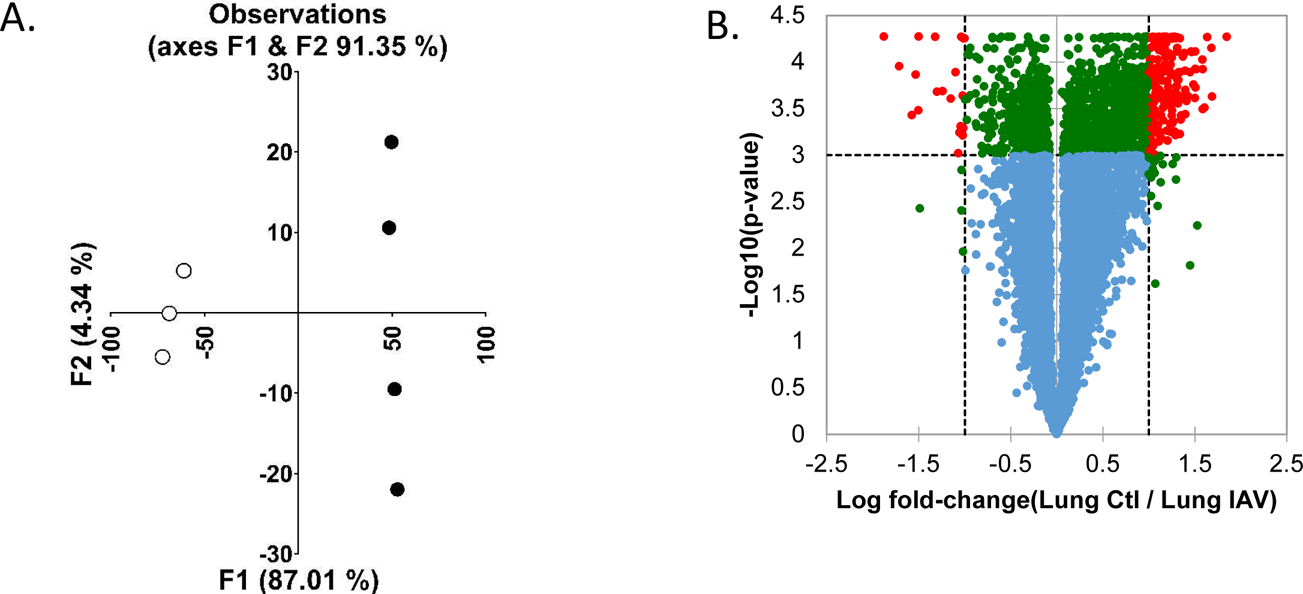
Unsupervised multivariant principal component analysis (PCA) of Tip DC and volcano plot of transcripts. At zero or nine days following intranasal infection of mice with IAV at (EID50: 2000) the lungs were homogenized and cells extracted. Cells were antibody stained for TipDC surface markers CD11b, Ly6c, GR-1, and MHCII. TipDC controls (Ctl) from day 0 were sorted based on high levels of CD11b, Ly6c, and GR-1. TipDC from day 9 of the IAV infection (IAV) were sorted based on high levels of CD11b, Ly6c, GR-1, and MHCII. cDNA libraries were generated from RNA, sequenced, and narrowed to confidentially identified transcripts. (A) Unsupervised multivariant principal component analysis (PCA) resulting in F1 and F2 with a cumulative percent variability of 78.56% Each circle represents a TipDC transcriptome, the open circles are control and the solid black IAV. (B) Differential expression differences were determined using Tukey HSD test for multiple comparisons with a Benjamini-Hochberg post-hoc false discover rate correction. 6642 transcripts were removed with nonspecific filtering to remove transcripts that were not modulated by IAV (i.e. 50% standard deviation threshold). Log10 p- values are plotted against the Log2 ratio

**S1 File.** Significantly enriched KEGG pathways altered by influenza infection of DC. PRIDE repository). (A) 7520 confidentially identified peptides from the stable isotopically labeled (SIL) and isobaric tagged (iTRAQ) proteomes and mapped to National Center for Biotechnology Information (NCBI) reference sequence accession numbers. (B&C) Confidently identified protein identifiers from the soluble and insoluble fractions were segregated into up regulated (B) or down regulated (C) based on 2-fold or greater abundances changes and submitted to DAVID for KEGG pathway analysis. Significantly enriched KEGG pathways are listed and gene symbols associated to NCBI accession numbers provided along with corresponding KEGG class and subclass from the BRITE hierarchy (colored circles correspond to subclasses from figure 1).

**S2 File.** TipDC transcriptome. At zero or nine days following intranasal infection of mice with IAV at (EID50: 2000) the lungs were homogenized and cells extracted. Cells were antibody stained for TipDC surface markers CD11b, Ly6c, GR-1, and MHCII. TipDC controls (Ctl) from day 0 were sorted based on high levels of CD11b, Ly6c, and GR-1. TipDC from day 9 of the IAV infection (IAV) were sorted based on high levels of CD11b, Ly6c, GR-1, and MHCII. cDNA libraries were generated from RNA, sequenced, and narrowed to confidentially identified transcripts. (A) Unsupervised multivariant principal component analysis (PCA) resulting in F1 and F2 with a cumulative percent variability of 78.56% Each circle represents a TipDC transcriptome, the open circles are control and the solid black IAV. (B) Differential expression differences were determined using Tukey HSD test for multiple comparisons with a Benjamini-Hochberg post-hoc false discover rate correction. (C) Transcripts that increased following IAV infection were submitted to DAVID and significantly enriched KEGG pathways are listed with corresponding KEGG class and subclass from the BRITE hierarchy. (D) Transcripts from the KEGG metabolic classes were submitted to DAVID and significantly enriched metabolic pathways are listed. (E) Transcripts that decreased following IAV infection were submitted to DAVID and significantly enriched KEGG pathways are listed with corresponding KEGG class and subclass from the BRITE hierarchy. (F) Unique transcripts from the Immune Pathway and Influenza A pathway were differentially expressed and corresponding protein names are listed under their functional category. (G) Gene symbols of the metabolic transcripts from TipDC and proteins from the in vitro differentiated DC are listed per pathway with the counts per cell type by down regulated (Down) or up regulate (Up) following IAV infection.

